# LAMBDA: A Prophage Detection Benchmark for Genomic Language Models

**DOI:** 10.64898/2026.03.26.714501

**Authors:** LeAnn M. Lindsey, Nicole L. Pershing, Keith Dufault-Thompson, Ho-jin Gwak, Anisa Habib, Aaron Schindler, Arjun Rakheja, June Round, W. Zac Stephens, Anne J. Blaschke, Hari Sundar, Xiaofang Jiang

## Abstract

Transformer-based genomic sequence models represent an emerging frontier in computational biology. Yet, their embeddings have not yet shown the same level of predictive power as natural and protein language models, highlighting a gap between current implementations and theoretical promise. Existing benchmarks for DNA language models primarily focus on classifying regulatory elements in eukaryotic genomes, leaving open the fundamental question of whether these models learn sequence-level features across whole genomes. We introduce LAMBDA, a benchmark designed to rigorously evaluate genome language model embeddings through phage-bacteria sequence discrimination across four categories of increasing complexity: probing tasks, fine-tuning assessments, diagnostic tests, and genome-wide prophage detection. Our comprehensive analysis of current genomic language models provides insight into the importance of training data selection relative to model size, the need for domain-specific training, and the capabilities and limitations of genomic language models for detecting prophage sequences. This benchmark represents a challenging genomic annotation task in the bacterial domain and addresses a key computational problem with direct relevance to microbiology and medicine.

## Introduction

Standardized benchmarks have been central to progress in the field of natural language processing. These benchmarks provide common frameworks for measuring performance, identifying model strengths and weaknesses, and guiding the development of increasingly capable models (49; 48; 25). Early benchmarks for genomic language models (gLMs) focused primarily on identifying short regulatory sequences in eukaryotic species, such as promoters (50), transcription factors (26; 52), and epigenetic marks (16). Two long-range benchmarks have recently been released (33; 12), which extend benchmark evaluations to longer sequence contexts, but remain focused on human regulatory genomics. While these benchmarks assess gLMs’ abilities to detect specific motifs and regulatory elements, they do not rigorously evaluate whether models can learn to identify functional boundaries or distinguish sequence types based on intrinsic properties, tasks that traditional methods accomplish through homology searches and explicit compositional features such as k-mer frequency distributions, GC content, and codon usage. Assessing whether gLMs can implicitly learn such patterns from sequence data alone would demonstrate a foundational understanding of DNA sequences with broad applicability in diverse genomic tasks.

In addition, several recent studies have questioned the “foundational” understanding of current genomic language models, demonstrating that, on current benchmarks, gLMs do not outperform randomly initialized models or simple supervised models for most genomic tasks (46; 45). These views stand in contrast to those of the model authors who claim that gLMs learn biologically meaningful representations of sequences, pointing to zero-shot performance across diverse tasks and the unsupervised discovery of genomic motifs as evidence of a foundational understanding (8; 30; 16; 52; 42).

Bacteriophages offer a compelling system in which to investigate these claims. Unlike model organisms with well-annotated genomes, the vast majority of phage genomes remain poorly characterized (21; 29). This makes them an ideal domain in which to test whether gLMs can identify meaningful sequence patterns, beyond what homology alone can reveal, a task with direct implications for the discovery of novel prophages.

Identifying the presence and location of prophages in host bacterial genomes is a significant goal for the microbiology and medical communities, but it remains a challenge (6). Phage genomes are extremely diverse, with closely related phage genomes often having a ‘mosaic’ nature (4; 17; 35), where homologous regions are separated by divergent or unrelated regions. Additionally, horizontal gene transfer in bacteria is commonly mediated by phages, which can lead to sections of a bacterial genome being integrated into the phage genome and then carried to different hosts. This process contributes to bacterial and phage diversity (10; 32; 24), but complicates the task of differentiating prophage and bacterial DNA. Lastly, phage genomes evolve rapidly, with high mutation rates and frequent gene loss, resulting in degraded prophage sequences being present throughout bacterial genomes (11). These factors result in a noisy genomic landscape in which distinguishing between phage and bacterial sequences is complex.

Currently, the most accurate prophage detection tools rely on sequence homology with known phage genes (51; 2; 38; 1; 27). Phage gene databases have grown in size and scope, but remain biased toward phages that infect common human pathogens, making it challenging for these reference-based methods to detect novel prophages, especially in less well-studied bacterial species (43). With advances in machine learning, sequence-based approaches that are not dependent on homology searches have become more common (28; 39). These approaches rely on feature engineering to distinguish between the two classes, using gene density, average gene length, differences in GC content, strand switching frequency, and percentages of specific start codons (22). These are often combined with searches for genes annotated with the term “phage” (40), which means that many of these methods still rely on sequence homology through gene annotation. One method employs a hybrid approach that integrates both homology features and sequence-based features into a deep learning classifier (9).

The Language Model Bacteriophage Detection Assessment (LAMBDA) evaluates foundational genomic understanding in DNA language models using bacteria and bacteriophage sequences as a model system (Figure 1). Our benchmark quantifies the impact of pretraining using probing experiments on pretrained embeddings compared to embeddings from randomly initialized models. It systematically investigates sources of prediction error, including compositional bias, false positive rates, and sensitivity across functional categories. The benchmark also includes genome-wide prophage detection tests against gold-standard datasets to critically assess whether the reported prophage detection capabilities of gLM models reflect a genuine biological signal and generalize outside of a limited set of well-studied bacterial hosts. In addition to providing a challenging and novel task for gLMs, prophage identification in bacterial genomes is of significant biological interest, with implications for the spread of antibiotic resistance and microbial evolution.

**Figure 1:**
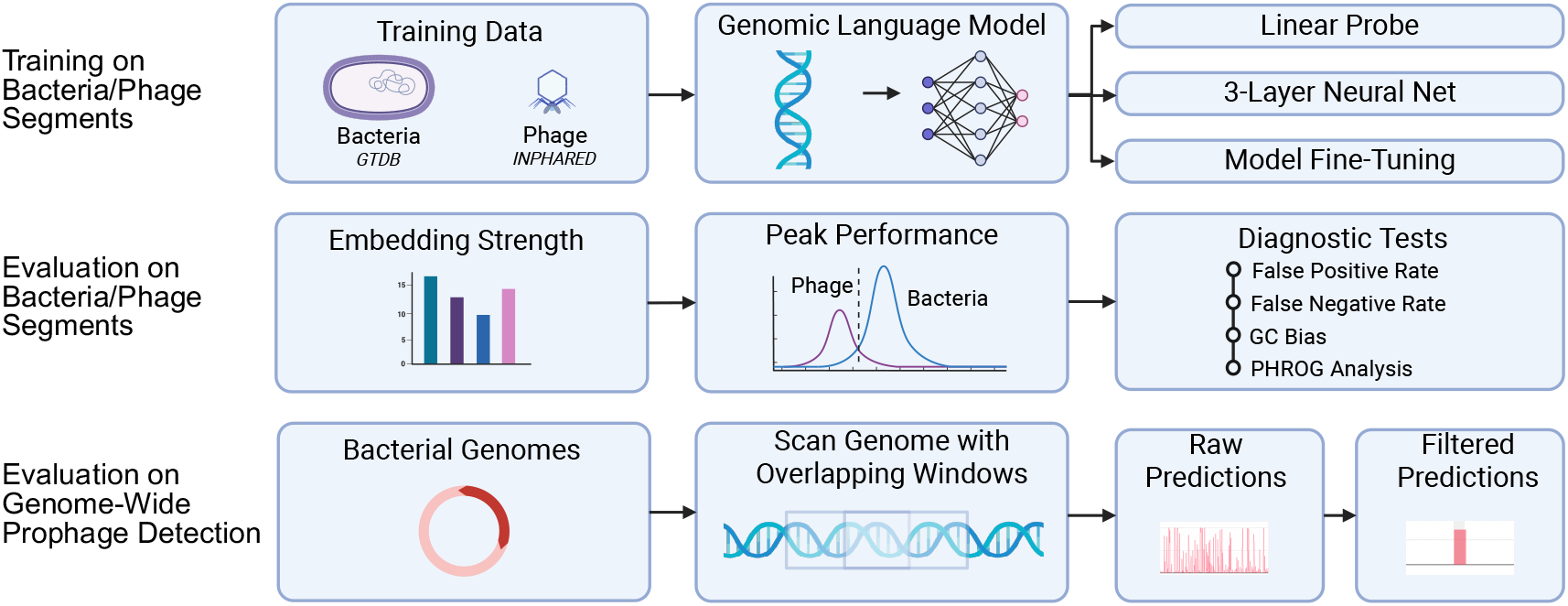
Overview of the LAMBDA benchmark. Models are trained on bacterial and phage genome fragments and evaluated using (i) embedding probes, (ii) model fine-tuning, (iii) diagnostic tests (GC bias, false positive/negative rates, and PHROG analysis), and (iv) genome-wide prophage detection on complete bacterial genomes.

## Materials and Methods

### Training dataset construction

#### Phage fragment selection

Complete phage genomes were obtained from the INPHARED (INfrastructure for a PHAge REference Database) database (14 Apr 2025 release) (14). Clusters with 95% average nucleotide identity (ANI) were generated using vclust (53), and one representative genome per cluster was selected, yielding 8,684 genomes for downstream processing. Gene annotations were obtained using Pharokka (7). Genomes were subsampled into fixed-length segments of 2000 nt, 4000 nt, and 8000 nt to create three datasets. Segments were sampled at random, non-overlapping positions at a rate of one segment per 10 kbp, with a minimum of one sample per genome.

#### Bacterial fragment selection

Bacterial genomes were selected from GTDB release r226 (36) representative genomes passing CheckM2 (37) quality filters (completeness ≥ 95%, contamination ≤ 5%), yielding 67,193 high-quality representatives; one genome per genus was retained (15,865 genomes) to prevent closely related genomes from appearing in multiple dataset partitions,, and genera were assigned to train, validation, and test in an 80:10:10 split with no genus shared across splits. Fragments of 2000, 4000, and 8000 nt were sampled from all contigs *≥* max(10,000 bp, 2 *×* fragment length), matched 1:1 to the phage fragment count in each split. To remove prophage-like sequence from the negatives, each bacterial fragment was aligned to the INPHARED phage-genome database with `blastn` (e-value ≤ 1*×*10^−5^, `max target seqs=2000`), and fragments with an alignment ≥ 200 bp at ≥ 70% nucleotide identity were removed. For each removed fragment, the source genome was aligned in full against INPHARED, the phage-matching regions were masked (padded *±*500 bp and merged per contig), and one replacement fragment was resampled from the unmasked regions of the same genome. This preserved every genome, its genus, and the exact 1:1 class balance while excluding prophage-derived sequence from the bacterial class. This screen is alignment-based against a reference phage database, and removes only prophage sequence homologous to known phages. Prophages highly divergent from all references may still be present, so the bacterial class is depleted of, but cannot be assumed to be entirely free of prophage sequences.

### Diagnostic dataset construction

#### GC-Control Dataset

Every sample in the test dataset had its nucleotide order shuffled to create a control dataset with sequences that preserved sequence length and GC content while destroying higher order sequence patterns.

#### Class Prediction Control Datasets

To estimate false positive rates on bacterial sequences, we created a bacteria-only benchmark consisting of 100 taxonomically balanced bacterial genomes drawn from the prophage-filtered GTDB test set. Genomes were selected to maximize taxonomic breadth. To estimate the false negative rates on phage sequences, we constructed a phage-only dataset from the test split (869 genomes).

#### PHROG Category Prediction Control Dataset

We created a second phage-only dataset using the 869 phage genomes in the test set, annotated with PHROG categories using Pharokka. In this PHROG category dataset, each sample was a coding sequence (CDS).

#### Genome-Wide Evaluation Set

To test the models against the challenge of scanning a full bacterial genome and detecting the presence of prophages, we assembled a reference dataset with published datasets that contain expert annotated prophage locations within bacterial genomes. We include the gold standard dataset published by Casjens et al. (11), which has been widely used by many prophage prediction programs, including PHASTER (2), as well as 16 additional prophage locations that were manually annotated by the authors of PHASTER. Another 62 reference locations were taken from Gauthier et al. (20), to add diversity. The final dataset spans 80 bacterial genomes representing 47 species across four phyla, which contain 380 annotated prophage locations. The viral taxonomy was obtained via Pharokka and identified 33 phage genera in two viral classes (Caudoviricetes and Faserviricetes), though the majority remain unclassified at the genus level. The reference bacterial accession numbers and the start and end positions for each predicted phage are provided (Supplementary Material Table S1).

### Evaluation Methods

#### valuation of genomic representations using frozen embeddings

To quantify the foundational strength of each model, two experiments were conducted. For each model, pretrained embeddings were extracted and used as fixed (frozen) representations to train two simple classifiers: a linear probe and a three-layer neural network classifier. A linear probe is a logistic regression classifier that maps the embeddings directly to the class predictions by fitting a hyperplane (a linear decision boundary) through the embedding space. This tests whether the two classes are linearly separable in the embedding representation. This approach, introduced by Alain (2016), has become a standard technique for evaluating the information encoded in different layers of a pre-trained model. The three-layer neural network classifier is a small feed-forward network with two hidden layers capable of measuring more complex, non-linear structure in the embeddings. We then extracted embeddings from each model using randomly initialized weights and used those embeddings as inputs in the same two classifiers. The metric used in this evaluation is the Matthews Correlation Coefficient (MCC) because it provides a balanced measure derived from all four entries of the confusion matrix and is robust to class imbalance. The model produces a score between +1 and *−*1, with a value of zero reflecting random chance. For each model, we report the difference in the Matthews Correlation Coefficient score, which quantifies the impact of model pretraining on the task.

#### Prophage Signal Extraction Algorithm

To reduce the number of false positive segments and increase precision, we used a filtering algorithm to consolidate the prophage signal from the genomic language models. We first collapsed the overlapping windows into a single score for each window using majority vote. After observing that the baseline level of prophage signal depends on the genome, we designed an algorithm that normalizes per-segment scores on a per-genome basis. We evaluated four normalization strategies, no normalization, z-score normalization, robust z-score normalization, and median subtraction, with z-score normalization having the most consistent results. Following normalization, we applied a bidirectional exponentially-weighted mean (EWM) smoothing. Normalized scores were then thresholded to produce a binary phage or non-phage call for each segment. Small gaps were merged into candidate prophage regions, and a length filter was applied to retain only candidates that exceeded the threshold length. To find the optimal hyperparameters for this algorithm, we conducted a grid search of all parameters separately for each model (Supplementary Material Figure S1, Table S2). The optimal hyperparameters were used for the final genome-wide evaluation scores.

#### Candidate Prophage Screening

To identify putative unannotated prophage regions, we scanned all 80 genomes for model predictions that do not overlap any ground truth region. Non-ground-truth predictions from all twenty models were pooled, and overlapping predictions across models were merged into consensus candidate regions. Candidates supported by two or more models were retained. Each candidate was flagged as “novel” if it was missed by both PHASTER and geNomad. Each candidate was manually annotated by extracting gene content from NCBI reference annotations and classifying each region as containing or lacking phage hallmark genes. “Confident phages” were defined as those with *≥* 10% structural genes with head/tail, “possible phages” with 5 *−* 10% structural, some evidence, “weak signal” was defined as having only 1-2 structural genes, likely degraded prophage. Those regions defined as “not phages” contained sequences that were either genomic islands, integrative conjugative elements (ICEs), or unlikely phages (Supplementary Material Table S3).

#### Models Evaluated

We evaluated ten genomic language models spanning a range of architectures, parameter counts, tokenization strategies, and pretraining corpora, alongside GC-content and k-mer frequency baselines and eight dedicated prophage-detection tools (Table 1). The models range from one million to seven billion parameters and were pretrained on corpora ranging from human and vertebrate genomes to prokaryotic and phage sequence, allowing us to assess the contribution of both scale and pretraining domain to prophage detection.

**Table 1:**
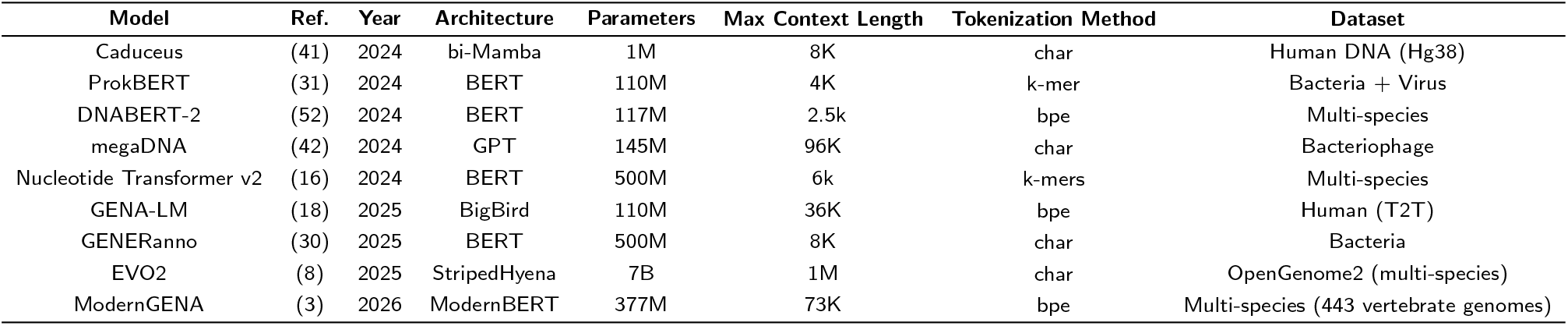
Genomic language models evaluated in the LAMBDA benchmark. For each model, we report the architecture, number of parameters, tokenization strategy, and training corpus. Models span a range of scales and pretraining regimes, including bacteria-specific, bacteriophage-specific, human DNA and multi-organism training datasets.

We compared the predictions from the genomic language models to seven traditional prophage detection tools and one protein language model-based tool. PIDE (19) is a recently published prophage detection tool that, to our knowledge, is the first published model built on protein language model embeddings. PHASTER is a widely used web server that identifies prophages through similarity to curated phage databases and structural feature annotation. We used the PHASTER Docker image for our screening. geNomad (9) integrates homology searches with a deep neural network classifier to detect viral sequences in microbial genomes. We also included VIBRANT (28), Phigaro (44), PhiSpy (1), PhageBoost (43), and VirSorter2 (22) in our comparisons. These tools collectively span homology-based, feature-based, and deep learning paradigms, providing a broad baseline for comparison.

## Results

### Evaluating the Power of Pretrained Sequence Representations

Across all models, pretrained embeddings outperformed randomly initialized embeddings under both linear probing and shallow nonlinear classification, though the margin varied (Supplementary Material Figure S2). At 8k context under linear probing, GENERanno and EVO2 both reached an MCC of 0.978, against random-embedding baselines of 0.410 and 0.687 respectively, while megaDNA improved from 0.284 to 0.903 and Nucleotide Transformer v2 from 0.575 to 0.945 (Figure 2, Supplementary Material Table S4). In contrast, DNABERT-2, Caduceus, and the GENA-LM family exhibited smaller gains from pretraining (ΔMCC *≤* 0.22) and correspondingly lower peak performance, indicating weaker representational power for this task. Notably, the nonlinear classifiers provided only modest gains over linear probes, indicating that with strong sequence representations, this task is largely linearly separable. Together, these results demonstrate that embedding strength is predictive, as models that show a small ΔMCC on this embedding test have weaker sequence representations and reduced downstream performance.

**Figure 2:**
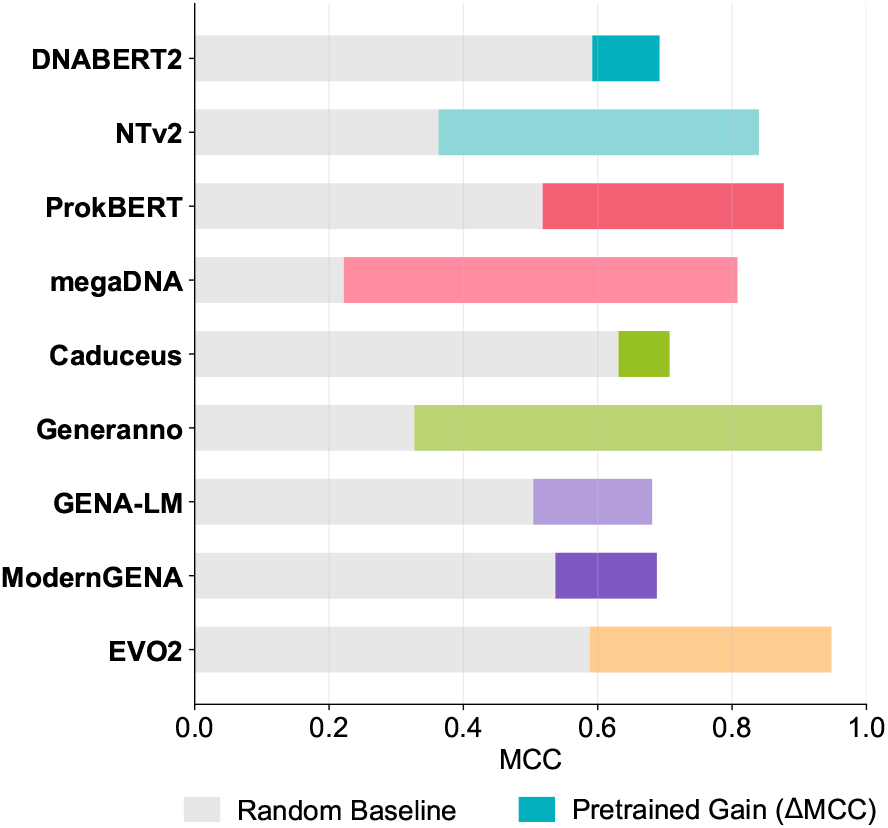
Embedding strength of genomic language models. Bars show the improvement in Matthews correlation coefficient (ΔMCC) obtained using pretrained embeddings compared to models with randomly initialized embeddings. Grey denotes random-baseline performance, and the colored portion represents the gain from pretraining. Higher ΔMCC values indicate stronger learned sequence representations.

### Evaluating Peak Model Performance

We define peak performance as a model’s best result across the linear probe, neural network, and fine-tuning experiments. At a matched input length of 2k, models differ substantially in peak performance, spanning a range from GENA-LM with an MCC score of 0.759 to EVO2 with an MCC score of 0.948. The ordering tracks the composition of each model’s pretraining corpus. Every model pretrained on prokaryotic or phage sequence reaches an MCC of at least 0.847 and every model pretrained predominantly on human or vertebrate genomes falls at or below 0.816 (Table 2). For reference, the GC-content baseline is only 0.16, whereas the k-mer frequency baseline is much stronger, with 6-mer counts reaching 0.80.

**Table 2:** Performance of genomic language models across fragment classification and genome-wide prophage detection in the LAMBDA benchmark. Fragment classification reports the Matthews correlation coefficient (MCC) on the held-out fragment test set for three readouts: a linear probe (LP), a 3-layer neural-network probe (NN), and full fine-tuning (FT; mean over five runs). Because the best fragment classifier did not always achieve the strongest genome-wide performance, all readouts were evaluated genome-wide, and the Model Used for Inference column identifies the readout used to generate the genome-wide predictions. Diagnostic metrics report GC-bias (MCC on GC-preserving shuffled sequences), the false positive rate (FPR) on bacteria-only genomes, and the false negative rate (FNR) on phage-only genomes. Genome-wide metrics report FPR, recall, and MCC across complete bacterial genomes before (Raw) and after (Filtered) post-processing. FPR, FNR, and recall are reported as fractions between 0 and 1. The best value in each genome-wide column is shown in bold.

| Model | Segment Length | Fragment Classification |  |  |  |  |  | Genome-wide Prophage Detection |  |  |  |  |  | Model Used for Inference |
| --- | --- | --- | --- | --- | --- | --- | --- | --- | --- | --- | --- | --- | --- | --- |
|  |  | MCC |  |  | Diagnostics |  |  | Raw |  |  | Filtered |  |  |  |
|  |  | LP | NN | FT | GC-bias | FPR | FNR | FPR | Recall | MCC | FPR | Recall | MCC |  |
| GC content | 2k | 0.157 | – | – | 0.157 | 0.411 | 0.417 | 0.375 | 0.391 | 0.015 | 0.069 | 0.295 | 0.124 | LR |
|  | 4k | 0.161 | – | – | 0.161 | 0.411 | 0.418 | 0.366 | 0.381 | 0.012 | 0.065 | 0.288 | 0.127 | LR |
|  | 8k | 0.142 | – | – | 0.142 | 0.413 | 0.413 | 0.364 | 0.382 | 0.012 | 0.094 | 0.337 | 0.128 | LR |
| k-mer $k = 4$ | 2k | 0.670 | 0.797 | – | 0.124 | 0.180 | 0.175 | 0.243 | 0.605 | 0.154 | 0.025 | 0.522 | 0.392 | LR |
|  | 4k | 0.739 | 0.867 | – | 0.119 | 0.156 | 0.149 | 0.172 | 0.569 | 0.211 | 0.037 | 0.582 | 0.404 | LR |
|  | 8k | 0.791 | 0.899 | – | 0.129 | 0.131 | 0.132 | 0.137 | 0.538 | 0.270 | 0.098 | 0.552 | 0.387 | LR |
| k-mer $k = 6$ | 2k | 0.727 | 0.802 | – | 0.092 | 0.159 | 0.139 | 0.240 | 0.643 | 0.170 | 0.021 | 0.548 | 0.430 | LR |
|  | 4k | 0.788 | 0.869 | – | 0.099 | 0.130 | 0.106 | 0.152 | 0.614 | 0.251 | 0.010 | 0.487 | 0.463 | LR |
|  | 8k | 0.854 | 0.918 | – | 0.061 | 0.106 | 0.082 | 0.111 | 0.593 | 0.329 | 0.052 | 0.536 | 0.473 | LR |
| DNABERT-2 | 2k | 0.692 | 0.752 | 0.770 | 0.132 | 0.105 | 0.119 | 0.187 | 0.581 | 0.184 | 0.012 | 0.525 | 0.477 | FT |
|  | 4k | 0.738 | 0.800 | 0.825 | 0.091 | 0.069 | 0.082 | 0.109 | 0.585 | 0.292 | 0.020 | 0.624 | 0.529 | FT |
| Nucleotide Transformer v2 | 2k | 0.840 | 0.874 | 0.864 | 0.057 | 0.064 | 0.080 | 0.148 | 0.723 | 0.324 | 0.007 | 0.687 | 0.662 | FT |
|  | 4k | 0.921 | 0.925 | 0.892 | (0.025) | 0.043 | 0.038 | 0.077 | 0.705 | 0.442 | **0.005** | 0.663 | 0.682 | NN |
|  | 8k | 0.945 | 0.958 | 0.934 | 0.036 | 0.030 | 0.023 | **0.037** | 0.634 | 0.515 | 0.009 | 0.668 | 0.671 | FT |
| Caduceus | 2k | 0.707 | 0.774 | 0.816 | 0.100 | 0.126 | 0.089 | 0.230 | 0.677 | 0.192 | 0.012 | 0.615 | 0.553 | FT |
|  | 4k | 0.774 | 0.843 | 0.877 | 0.159 | 0.073 | 0.079 | 0.115 | 0.613 | 0.307 | 0.013 | 0.656 | 0.601 | FT |
|  | 8k | 0.833 | 0.870 | 0.901 | 0.128 | 0.064 | 0.098 | 0.096 | 0.609 | 0.365 | 0.011 | 0.601 | 0.574 | FT |
| ProkBERT-mini | 2k | 0.825 | 0.857 | 0.870 | 0.103 | 0.078 | 0.069 | 0.132 | 0.661 | 0.294 | 0.007 | 0.689 | 0.667 | NN |
| ProkBERT-mini-c | 2k | 0.807 | 0.845 | 0.897 | 0.120 | 0.077 | 0.048 | 0.146 | 0.662 | 0.277 | 0.005 | 0.614 | 0.628 | NN |
| ProkBERT-mini-long | 2k | 0.877 | 0.917 | 0.895 | (0.002) | 0.047 | 0.051 | 0.116 | 0.745 | 0.380 | 0.007 | 0.713 | 0.704 | NN |
| megaDNA | 2k | 0.808 | 0.847 | – | 0.033 | 0.070 | 0.099 | 0.137 | 0.650 | 0.271 | 0.006 | 0.676 | 0.657 | NN |
|  | 4k | 0.871 | 0.904 | – | (0.021) | 0.050 | 0.059 | 0.089 | 0.652 | 0.395 | 0.006 | 0.652 | 0.651 | NN |
|  | 8k | 0.903 | 0.933 | – | 0.049 | 0.034 | 0.084 | 0.071 | 0.628 | 0.487 | 0.007 | 0.634 | 0.633 | NN |
| GENERanno | 2k | 0.934 | 0.928 | 0.897 | 0.079 | 0.043 | 0.035 | 0.179 | 0.792 | 0.359 | 0.009 | 0.698 | 0.649 | FT |
|  | 4k | 0.965 | 0.966 | 0.927 | 0.098 | 0.026 | 0.012 | 0.426 | 0.822 | 0.290 | 0.029 | 0.606 | 0.475 | NN |
|  | 8k | **0.978** | **0.981** | **0.945** | 0.145 | **0.024** | **0.006** | 0.321 | 0.759 | 0.366 | 0.030 | 0.649 | 0.532 | FT |
| GENA-LM | 2k | 0.681 | 0.759 | 0.657 | 0.137 | 0.126 | 0.131 | 0.247 | 0.587 | 0.140 | 0.010 | 0.495 | 0.470 | NN |
|  | 4k | 0.749 | 0.842 | 0.701 | 0.099 | 0.025 | 0.525 | 0.109 | 0.423 | 0.168 | 0.008 | 0.395 | 0.399 | NN |
|  | 8k | 0.802 | 0.873 | 0.739 | 0.144 | 0.068 | 0.081 | 0.123 | 0.538 | 0.269 | 0.018 | 0.488 | 0.499 | NN |
| ModernGENA | 2k | 0.688 | 0.782 | 0.804 | 0.138 | 0.102 | 0.110 | 0.210 | 0.632 | 0.201 | 0.011 | 0.584 | 0.529 | FT |
|  | 4k | 0.759 | 0.859 | 0.839 | 0.129 | 0.095 | 0.073 | 0.076 | 0.506 | 0.339 | 0.015 | 0.612 | 0.540 | FT |
|  | 8k | 0.813 | 0.889 | 0.914 | 0.106 | 0.035 | 0.050 | 0.068 | 0.598 | 0.448 | 0.015 | 0.638 | 0.613 | FT |
| EVO2 | 2k | 0.948 | 0.940 | – | **0.001** | 0.055 | 0.029 | 0.111 | 0.782 | 0.479 | 0.005 | 0.719 | **0.722** | LP |
|  | 4k | 0.966 | 0.963 | – | (0.121) | 0.043 | 0.018 | 0.090 | 0.731 | 0.534 | 0.010 | **0.767** | 0.716 | LP |
|  | 8k | **0.978** | 0.970 | – | (0.042) | 0.032 | 0.019 | 0.086 | 0.666 | **0.543** | 0.011 | 0.730 | 0.688 | LP |
| EVO2 + SAE | 2k | – | – | – | – | – | – | 0.274 | **0.888** | 0.301 | 0.010 | 0.755 | 0.686 | SAE |
|  | 4k | – | – | – | – | – | – | 0.226 | 0.880 | 0.354 | 0.012 | 0.757 | 0.679 | SAE |
|  | 8k | – | – | – | – | – | – | 0.226 | 0.872 | 0.380 | 0.013 | 0.733 | 0.658 | SAE |

“–” denotes not applicable (baselines and EVO2/megaDNA were not fine-tuned, and GC content has no 3-layer NN). For the feature baselines the deployed model is logistic regression (LR). GC-bias values in parentheses are negative.

### Evaluating Sources of Prediction Error

#### Testing for GC-composition bias

On the GC-composition bias test, all models had MCC values that fell within ±0.16 of zero, indicating that GC-composition accounts for only a small fraction of model performance. GC-bias was most pronounced for ModernGENA (0.138), GENA-LM (0.137), and DNABERT-2 (0.132), and smallest for EVO2 (0.001) and ProkBERT-mini-long (−0.002), reported at the 2k segment length.

#### Testing for Class Prediction Bias

On the balanced control datasets at 2k, error rates were low and approximately symmetric. False positives on bacteria-only sequences averaged 8.1% and false negatives on phage-only sequences averaged 7.8% (Table 2). Genome-wide false-positive rates were approximately twofold higher (17.6% against 8.1% at 2k), a gap that holds for every model tested, indicating that balanced fragment classification understates the error encountered when scanning complete genomes.

#### Testing Performance Across PHROG Functional Categories

Sensitivity varied considerably across both categories and models (Table 3). Each phage coding sequence in this evaluation was assigned to a functional category from the PHROG database, which clusters phage proteins by remote homology into classes spanning the virion structural machinery (Head & Packaging, Tail, Connector), genome replication (DNA, RNA & Nucleotide Metabolism), host lysis (Lysis), lysogenic control (Integration & Excision, Transcription Regulation), and host manipulation (Moron, AMG & Host Takeover). The most reliably detected categories were Head & Packaging, with an average sensitivity of 0.76, and Tail proteins, comprising the fiber and baseplate proteins that mediate host recognition with an average sensitivity of 0.74. Uncharacterized genes were detected least reliably, with a sensitivity of 0.60 for those with a PHROG match but no known function (27.8% of genes), and 0.45 for those with no PHROG match at all (38.7% of genes). These two categories together comprise 66.5% of the annotated genes in this dataset, whereas the two best-detected categories account for only 14.1%.

**Table 3:** Phage gene detection sensitivity stratified by PHROG functional categories across genomic language models. PHROG (Phage Orthologous Groups) categories represent curated functional groupings of phage genes based on sequence homology. Evaluation is performed on a fully annotated phage dataset, where each gene is assigned to a PHROG category. “% of Total” indicates the fraction of annotated phage genes belonging to each category in this dataset. Sensitivity is defined as the fraction of annotated phage genes in each category that the model correctly identifies as phage. Categories are ordered by average sensitivity across models (highest to lowest).

| Category | % of Total | GC | k-mer4 | k-mer6 | DNABERT-2 | NTv2 | ProkBERT | megaDNA | Caduceus | GENERanno | GENA-LM | ModernGENA | EVO2 | EVO2+SAE | Avg Sensitivity |
| --- | --- | --- | --- | --- | --- | --- | --- | --- | --- | --- | --- | --- | --- | --- | --- |
| Head & Packaging | 6.7% | 0.54 | 0.86 | 0.83 | 0.83 | 0.84 | 0.70 | 0.33 | 0.41 | 0.87 | 0.92 | 0.83 | 0.99 | 0.89 | 0.76 |
| Tail | 7.4% | 0.51 | 0.82 | 0.80 | 0.78 | 0.82 | 0.69 | 0.37 | 0.39 | 0.83 | 0.90 | 0.79 | 0.98 | 0.88 | 0.74 |
| DNA, RNA & Nucleotide Metabolism | 9.8% | 0.59 | 0.83 | 0.81 | 0.81 | 0.76 | 0.61 | 0.28 | 0.28 | 0.82 | 0.92 | 0.79 | 0.96 | 0.90 | 0.72 |
| Other | 3.0% | 0.64 | 0.80 | 0.78 | 0.74 | 0.71 | 0.44 | 0.22 | 0.15 | 0.68 | 0.90 | 0.71 | 0.91 | 0.83 | 0.66 |
| Lysis | 1.8% | 0.50 | 0.78 | 0.76 | 0.66 | 0.65 | 0.44 | 0.24 | 0.16 | 0.65 | 0.88 | 0.69 | 0.96 | 0.92 | 0.64 |
| Connector | 1.5% | 0.44 | 0.78 | 0.74 | 0.72 | 0.62 | 0.42 | 0.18 | 0.08 | 0.71 | 0.92 | 0.69 | 0.99 | 0.95 | 0.63 |
| Moron, AMG & Host Takeover | 1.4% | 0.68 | 0.79 | 0.75 | 0.65 | 0.64 | 0.36 | 0.19 | 0.13 | 0.63 | 0.86 | 0.66 | 0.89 | 0.69 | 0.61 |
| Unknown Function<sup>†</sup> | 27.8% | 0.61 | 0.78 | 0.74 | 0.57 | 0.56 | 0.37 | 0.15 | 0.08 | 0.67 | 0.89 | 0.64 | 0.95 | 0.81 | 0.60 |
| Integration & Excision | 0.5% | 0.47 | 0.65 | 0.67 | 0.64 | 0.57 | 0.48 | 0.16 | 0.23 | 0.48 | 0.82 | 0.62 | 0.92 | 0.88 | 0.58 |
| Transcription Regulation | 1.3% | 0.67 | 0.71 | 0.68 | 0.61 | 0.58 | 0.29 | 0.11 | 0.02 | 0.58 | 0.86 | 0.64 | 0.94 | 0.85 | 0.58 |
| No PHROG<sup>‡</sup> | 38.7% | 0.58 | 0.66 | 0.62 | 0.30 | 0.32 | 0.20 | 0.10 | 0.02 | 0.40 | 0.73 | 0.41 | 0.89 | 0.60 | 0.45 |
| **Overall** | **100%** | **0.58** | **0.75** | **0.71** | **0.53** | **0.53** | **0.38** | **0.18** | **0.13** | **0.61** | **0.83** | **0.59** | **0.93** | **0.75** | **0.58** |

<sup>†</sup>Matched to a PHROG entry but no known biological function assigned
<sup>‡</sup>No match to any entry in the PHROG database

Model performance on CDS-length segments diverged sharply from performance on fixed 2k fragments. Caduceus (0.13 overall) and megaDNA (0.18) detected almost no phage CDS despite each missing under 10% of phage sequences in the fragment-level control, indicating severe degradation on short inputs. EVO2 was the most robust with a sensitivity of 0.93. The k-mer baseline had very strong performance on this task, with 4-mer counts (0.75) exceeding every model except EVO2 and GENA-LM.

### Genome-Wide Prophage Detection

#### Extracting Prophage Signal from Raw Model Predictions

The genome-wide prophage detection task is considerably more challenging for genomic language models than controlled segment classification. The false positive rate increased across all models relative to the controlled experiments. Per-genome normalization and filtering substantially reduced false positives, increasing MCC for every model at every input length (mean +0.26) while decreasing recall only slightly (mean *−*3.3 percentage points) (Figure 3). After filtering, EVO2 achieved the highest overall performance (MCC 0.722), followed by ProkBERT-mini-long (0.704), EVO2+SAE (0.686), and Nucleotide Transformer v2 (0.682). GENA-LM (0.499) and DNABERT-2 (0.529) had the lowest performance.

**Figure 3:**
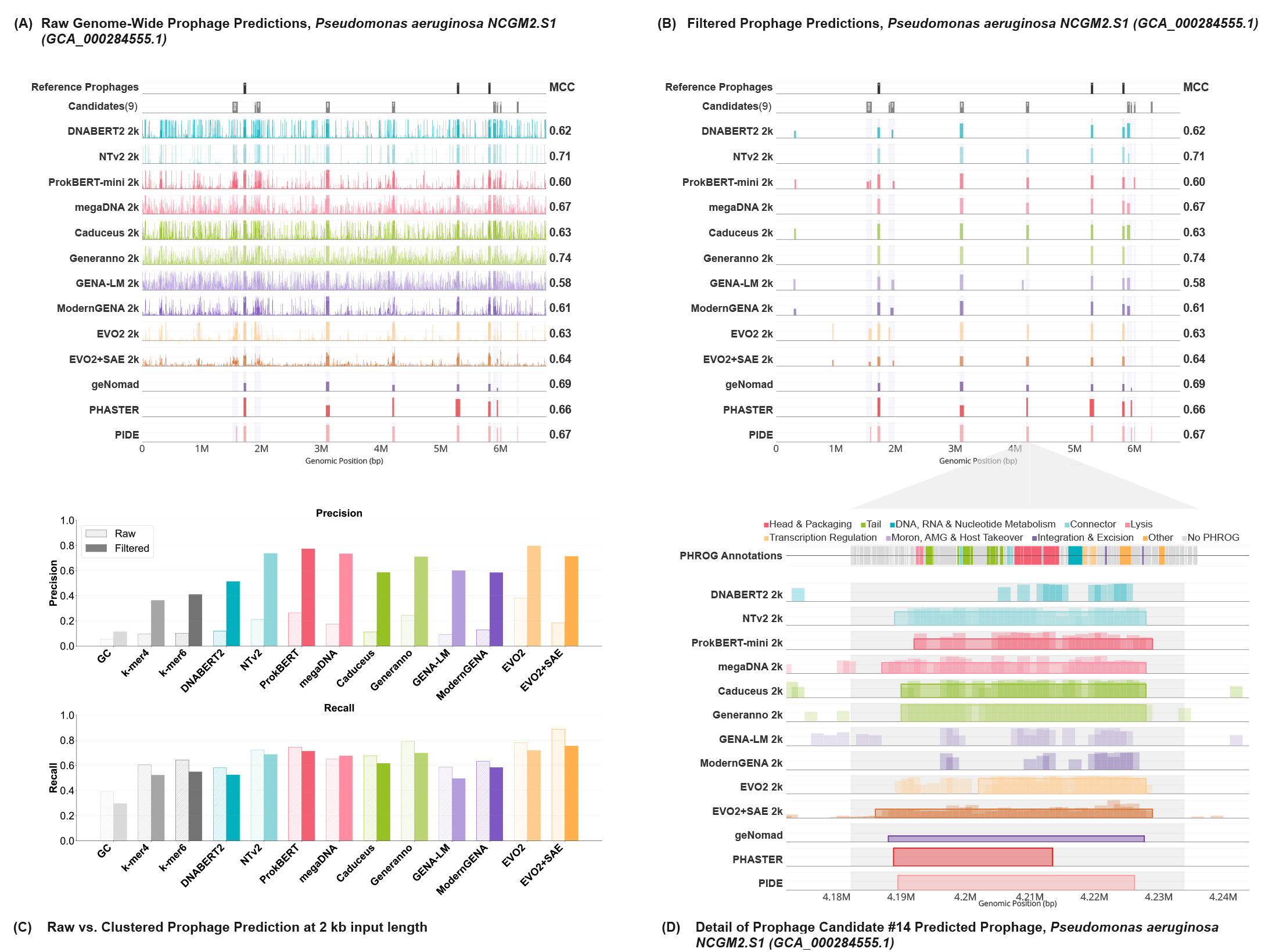
Genome-wide prophage detection and post-processing in LAMBDA. (A) Raw genome-wide predictions for a representative bacterial genome (Pseudomonas aeruginosa NCGM2.S1), showing the per-window prophage scores produced by each model before post-processing. Black bars indicate published reference prophage regions, while the “Candidates” track shows regions supported by two or more models that are absent from the reference annotations. (B) Predictions after per-genome normalization, smoothing, merging adjacent high-scoring windows, and length filtering to generate final prophage calls. (C) Genome-wide performance before (Raw) and after post-processing (Filtered), showing improved precision and MCC with minimal loss of recall. (D) Detailed view of candidate prophage #14, comparing predictions from genomic language models, traditional prophage detection tools, and the protein language model PIDE with PHROG functional annotations.

In contrast to the segment-level tests, where every model improved with additional context, longer context did not consistently improve genome-wide performance, with half of the models scoring lower at 8k than at 2k, including EVO2, the highest performer overall. Per-genome performance varied widely across the benchmark, from an average gLM MCC of 0.945 in *Mycobacteroides abscessus* GD17 to 0.198 in *Escherichia coli* K-12 (Supplementary Material Figures S3-S6). Detection performance varied widely with both host taxonomy and viral lineage, with median MCC ranging from 0.85 in Actinomycetota to 0.59 in Pseudomonadota at the phylum level (Supplementary Material Figures S7-S11). Detection quality also declined as prophages diverged from previously characterized phages: the overlap between predicted and reference regions (Jaccard index) fell steadily with increasing MASH distance to the nearest known phage (Pearson *r* = −0.51, *p* < 10^−17^) (Supplementary Material Figures S12-S13).

The compositional baselines, which were competitive with the models on fragment classification, separated clearly from the gLMs on the genome-wide task. The strongest baseline (6-mer frequencies) reached a filtered MCC of only 0.473, below every genomic language model, while GC content alone reached only 0.128 (Table 2).

#### Scanning for Novel, Unannotated Prophage Regions

While defining prophage boundaries is inherently difficult, our scanning presented an opportunity to potentially detect novel, unannotated phage regions. Across the 80 bacterial genomes in our benchmark, we identified 305 candidate regions that were supported by between two and twenty models (median: four) that did not overlap with any annotation in the published ground-truth dataset. To characterize these candidates, we performed a manual review of all 305 regions, evaluating each candidate for structural-gene, integration/excision, lysis, and annotation evidence (Supplementary Material Table S3). This review yielded 22 candidates classified as *likely phage*, 92 as *possible phage*, 113 as *weak phage signal*, 31 as *genomic island / ICE*, 16 as *genomic island*, and 31 as *unlikely phage*. Even within the 36 high-confidence candidates supported by at least ten models, only nine were classified as likely phage, whereas fourteen showed weak phage signal, eleven were classified as possible phage, one as unlikely phage, and one as a genomic island / ICE. A representative strong example within this set was a ∼44 kb region in *Gordonia iterans* Co17 (candidate 39), supported by 17 models and classified as likely phage with eight head/packaging, ten tail, three connector, and three lysis genes.

For comparison, the 380 reference prophage regions were supported by a median of 14 of the 20 detection methods, with 351 (92.4%) supported by at least two, the same criterion applied to the candidate regions. Likely-phage candidates were supported by a median of eight methods and the remaining candidates by a median of three (Supplementary Material Figure S14). Thirteen reference regions received no support from any method, and these are the prophages least represented in the phage training set (Supplementary Material Figure S15).

Across all 305 candidates, the predominant pattern was therefore ambiguous or non-phage mobile elements rather than intact prophages: 191 regions were classified as weak phage signal, genomic island/ICE, genomic island, or unlikely phage. These results indicate that most candidate regions flagged outside the ground truth are not spurious predictions but rather reflect genuine mobile genetic elements whose compositional and structural signatures overlap with those of prophages. This overlap between prophages and other mobile elements represents a fundamental challenge for all prophage detection methods and is a likely source of false-positive predictions across both gLMs and traditional tools. At the same time, the presence of 22 likely phage candidates absent from the published annotations suggests that the ground-truth dataset remains incomplete, and that the MCC scores we report are correspondingly conservative.

#### Probing EVO2 Latent Space with Sparse Autoencoders

Sparse autoencoders (SAEs) are mechanistic interpretability tools that decompose the internal representations of a language model into a sparse set of interpretable features (15; 34). The recently published EVO2 model reports prophage detection as one of several biological features that EVO2 has learned using unsupervised learning. In their study, a sparse autoencoder (SAE) is used to extract information from the model’s internal neurons. One of the identified features, f/19746, was associated with prophage sequences across the prokaryotic domain. We used the EVO2 model, along with the trained SAE, to extract this signal for all genomes in our test set to evaluate its generalizability as a prophage detection tool.

We found that f/19746 activation varies considerably across genomes, firing cleanly in prophage regions of some genomes, but producing noisy baseline signal or sparse, non-prophage-associated activations in others. After applying our clustering algorithm, EVO2+SAE achieved competitive but lower performance than the EVO2 neural network classifier (average MCC of 0.674 vs. 0.709) (Supplementary Material Figures S16-S18).

### Comparison with Traditional and Protein-based Models

A comparison with traditional prophage detection tools shows that genomic language model performance on this task lags slightly behind traditional methods and is comparable with PIDE, a model built using protein language model embeddings (Table 4). EVO2 had the highest performance of the gLMs, with an MCC of 0.719, but this score was 0.075 lower than the model with the highest performance (geNomad). The gap is driven by recall rather than precision. EVO2’s precision (0.780) is the second highest of any method evaluated and exceeds geNomad’s (0.761), but its recall (0.718) trails geNomad’s (0.873). No gLM reached the recall of the four leading traditional methods, whereas several matched or exceeded their precision.

**Table 4:**
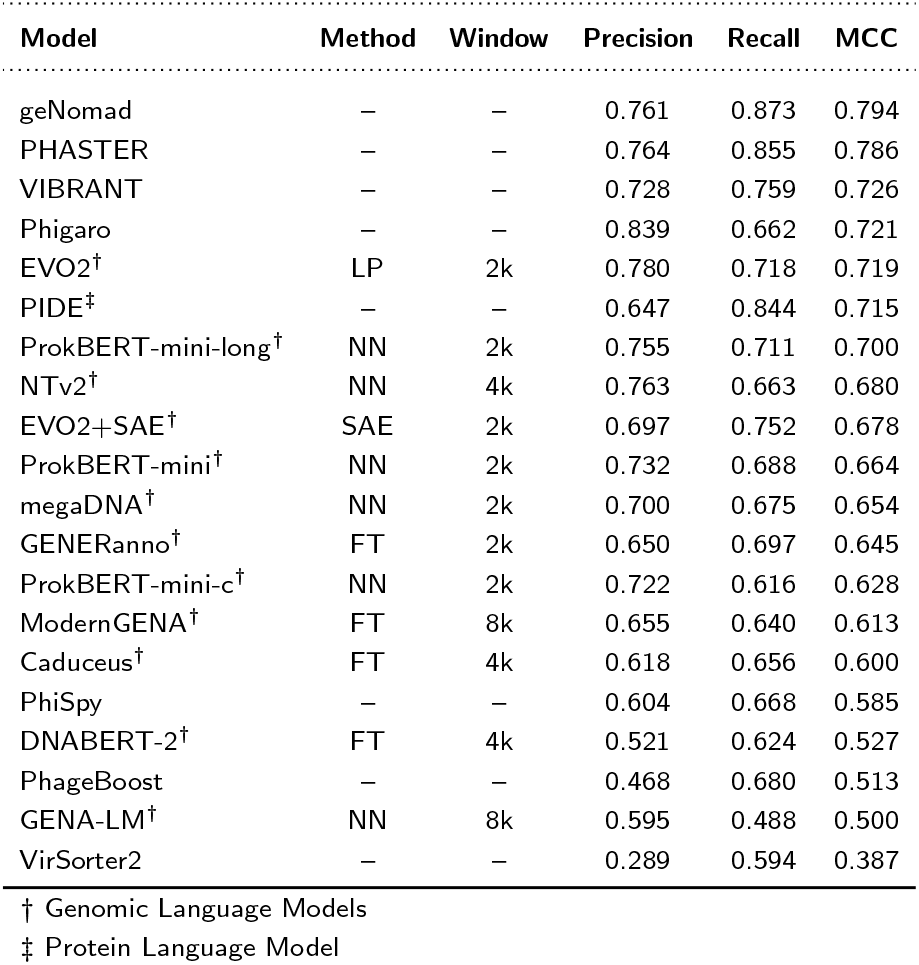
Comparison of genomic language models with traditional prophage detection tools. Models are evaluated on genome-wide prophage detection and ranked by Matthews Correlation Coefficient (MCC). For each genomic language model the best-of configuration is shown (deployment head and fragment window that maximize genome-wide MCC); the Method and Window columns record that choice. Shaded rows indicate genomic language models (gLMs) fine-tuned for prophage detection a protein language model-based approach is included for reference. Traditional tools represent current state-of-the-art methods for prophage identification.

## Discussion

Standardized benchmarks that replicate unique domain challenges provide a foundation for the development and evaluation of new models. The LAMBDA benchmark provides this for DNA language models, presenting novel tasks that represent the unique challenges of detecting prokaryotic and viral sequence-level features in DNA. Our results demonstrate that previous studies showing randomly initialized models having similar or better performance than pretrained genomic language models (47) likely reflect insufficient benchmark difficulty rather than a lack of representational benefit from pretraining. The LAMBDA probing tasks quantify the contribution of pretraining and show that across all models tested, pretrained embeddings significantly outperform their randomly initialized counterparts. This demonstrates that benchmark rigor is essential for evaluating the true sequence representational quality of genomic language models.

Published genomic language models differ simultaneously in architecture, scale, tokenization, and pretraining data composition, making it difficult to isolate the contribution of any individual factor. Within this benchmark, however, pretraining data composition tracked performance more closely than scale did. Although EVO2, with seven billion parameters, achieved the highest performance on the genome-wide prophage detection task, the second-highest was ProkBERT-mini-long (MCC 0.700), which has roughly sixty-fold fewer parameters but was pretrained on a curated prokaryotic dataset. This pattern held across the benchmark, where the highest-performing models were consistently those pretrained on corpora containing prokaryotic or phage sequence, spanning parameter counts from 110 million to seven billion. Within this group, model size was a poor predictor of relative performance. This has significant implications for the development of future gLMs, where the relative contributions of model size and training data curation need to be weighed against the computational and human costs of developing new models. As our results demonstrate, some tasks may require smaller, domain-specific models to achieve consistent and high performance.

These results demonstrate that the domain and task alignment are stronger predictors of model performance than model scale. This has significant implications for the development of future gLMs, where the relative contributions of model size and training data curation need to be weighed against the computational and human costs of developing new models.

The compositional baselines were competitive with genomic language models on fragment classification, but fell below every model genome-wide, suggesting that resolving prophage boundaries within an intact bacterial genome requires higher-order sequence features that the genomic sequence models capture and simple composition does not.

Defining a complete ground truth for prophage content in bacterial genomes is inherently difficult. The annotation of prophages depends on the tools and reference databases available at the time of curation. Further complicating this task is the fact that the boundaries between prophages, integrative conjugative elements (ICEs), pathogenicity islands, and other mobile genetic elements are not always clear-cut, as these elements often share core integration and mobilization machinery. This overlap between prophages and other mobile elements poses a fundamental challenge for all prophage detection methods and is likely a source of false-positive predictions in both gLMs and traditional tools. At the same time, the detection of nearly complete prophages in the *Y. pestis* genomes demonstrates that the ground-truth dataset itself is incomplete, needing additional experimental validation to identify and characterize phages. This also implies that the MCC scores that we report are likely conservative, being impacted by the incomplete knowledge of phages in bacterial genomes.

Our analysis also provides an opportunity to evaluate the potential impact of new approaches within the field. For example, our analysis of the EVO2 sparse autoencoder output indicated that SAEs represent a promising avenue for future research, but that the EVO2+SAE feature f/19746 does not generalize uniformly across the prokaryotic domain. We propose two possible explanations for this limited generalizability, insufficient domain-specific training data and distributed encoding of the prophage signal. The extracted feature can only reflect what the model has learned during training, and since a large portion of the EVO2 dataset was focused on regulatory information in the eukaryotic domain, the model may be undertrained on specific bacterial taxa. Second, the prophage signal may not be fully encoded in a single feature from one layer but could be distributed across a circuit of interacting neurons, only one of which has been identified. Circuit-level representations have been demonstrated in transformer language models (5; 13; 15; 23), suggesting that a more comprehensive circuit-level analysis may be needed to capture the full prophage signal across the entire prokaryotic domain.

## Supporting information

LAMBDA v1 Supplementary Figures

LAMBDA v1 Supplementary Tables

## Limitations

Although we used the batch sizes and learning rates recommended by the authors for each model, the potential benefits of extensive hyperparameter tuning for model performance remain to be explored. Similarly, signal extraction methods can significantly affect final performance metrics, and more research is needed to identify the most effective and consistent approaches. Additionally, as our manual review of 305 candidate regions demonstrates, the boundaries between prophages, ICEs, and other mobile genetic elements are often ambiguous, and the ground-truth dataset likely remains incomplete despite expert curation. This overlap between prophage and non-phage mobile element signatures represents a fundamental challenge for all detection methods and a source of both false-positive and false-negative predictions. Further analyses, including systematic comparisons with mobile genetic element datasets, are needed to more fully characterize gLM prediction limitations and inform future improvement strategies.

## Conclusion

We conclude that current genomic language models capture meaningful DNA sequence representations relevant to the prophage detection annotation task, although homology-based tools still achieve higher performance on this task. LAMBDA addresses a critical gap in existing benchmarks by providing specific metrics that reveal where models fail and moves beyond classification tasks toward genome-wide screening for specific sequence patterns and motifs. This rigorous framework highlights the current limitations of genomic language models. The consistent relationship we observed between the representational power of embeddings and downstream accuracy suggests that improvements in pretraining and training data curation will translate into meaningful gains on genomic annotation capabilities.

## Acknowledgments

We thank Yair Schiff, Zhihan Zhou, Brien Hie and Eric Nguyen for advice and model specific recommendations. We thank Gaetano Manzo, Antonio Pedro Camargo, Aditya Bhaskara, Saday Sadayappan and Henry Secaira for helpful discussions and advice. This work utilized the computational resources of the NIH HPC Biowulf cluster (https://hpc.nih.gov), computational resources and support from the Center for High Performance Computing at the University of Utah as well as Bridges-2 at Pittsburgh Supercomputing Center and Delta at the National Center for Supercomputing Applications (NCSA) through allocation BIO230092 from the Advanced Cyberinfrastructure Coordination Ecosystem: Services & Support (ACCESS) program, which is supported by National Science Foundation grants #2138259, #2138286, #2138307, #2137603, and #2138296.

## Author contributions statement Data Availability

All datasets used in this study were derived from publicly available sources. Bacterial genomes were obtained from the Genome Taxonomy Database (GTDB), and phage genomes were obtained from the INfrastructure for a PHAge REference Database (INPHARED).

The processed datasets used in the LAMBDA benchmark, including training, evaluation, and diagnostic test sets are available for download from https://doi.org/10.5281/zenodo.19236553.

## Code Availability

All code required to reproduce the LAMBDA benchmark, including model training, evaluation, and analysis pipelines, is available on GitHub at https://github.com/leannmlindsey/LAMBDA and archived on Zenodo at https://doi.org/10.5281/zenodo.21909098.

An interactive web-based visualization tool is available to visualize the genome-wide prophage predictions on the test set, including results from the EVO2 + sparse autoencoder (SAE) analysis. https://leannmlindsey.github.io/lambda-benchmark/.

## References

1. Sajia Akhter, Ramy K. Aziz, and Robert A. Edwards. PhiSpy: a novel algorithm for finding prophages in bacterial genomes that combines similarity- and composition-based strategies. Nucleic Acids Research, 40(16):e126, September 2012.

2. David Arndt, Jason R. Grant, Ana Marcu, Tanvir Sajed, Allison Pon, Yongjie Liang, and David S. Wishart. PHASTER: a better, faster version of the PHAST phage search tool. Nucleic Acids Research, 44(Web Server issue):W16–W21, July 2016.

3. Alena Aspidova, Yuri Kuratov, Artem Shadskiy, Mikhail Burtsev, and Veniamin Fishman. Back to BERT in 2026:ModernGENA as a Strong, Efficient Baseline for DNA Foundation Models, April 2026. ISSN: 2692-8205 Pages: 2026.04.21.719816 Section: New Results.

4. Mahdi Belcaid, Anne Bergeron, and Guylaine Poisson. Mosaic graphs and comparative genomics in phage communities. Journal of Computational Biology, 17(9):1315–1326, September 2010.

5. Adithya Bhaskar, Alexander Wettig, Dan Friedman, and Danqi Chen. Finding Transformer Circuits with Edge Pruning, April 2025. arXiv:2406.16778 [cs].

6. Ho Bin Jang, Benjamin Bolduc, Olivier Zablocki, Jens H. Kuhn, Simon Roux, Evelien M. Adriaenssens, J. Rodney Brister, Andrew M. Kropinski, Mart Krupovic, Rob Lavigne, Dann Turner, and Matthew B. Sullivan. Taxonomic assignment of uncultivated prokaryotic virus genomes is enabled by gene-sharing networks. Nature Biotechnology, 37(6):632–639, June 2019.

7. George Bouras, Roshan Nepal, Ghais Houtak, Alkis James Psaltis, Peter-John Wormald, and Sarah Vreugde. Pharokka: a fast scalable bacteriophage annotation tool. Bioinformatics, 39(1):btac776, January 2023.

8. Garyk Brixi, Matthew G. Durrant, Jerome Ku, Michael Poli, Greg Brockman, Daniel Chang, Gabriel A. Gonzalez, Samuel H. King, David B. Li, Aditi T. Merchant, Mohsen Naghipourfar, Eric Nguyen, Chiara Ricci-Tam, David W. Romero, Gwanggyu Sun, Ali Taghibakshi, Anton Vorontsov, Brandon Yang, Myra Deng, Liv Gorton, Nam Nguyen, Nicholas K. Wang, Etowah Adams, Stephen A. Baccus, Steven Dillmann, Stefano Ermon, Daniel Guo, Rajesh Ilango, Ken Janik, Amy X. Lu, Reshma Mehta, Mohammad R. K. Mofrad, Madelena Y. Ng, Jaspreet Pannu, Christopher Ré, Jonathan C. Schmok, John St John, Jeremy Sullivan, Kevin Zhu, Greg Zynda, Daniel Balsam, Patrick Collison, Anthony B. Costa, Tina Hernandez-Boussard, Eric Ho, Ming-Yu Liu, Thomas McGrath, Kimberly Powell, Dave P. Burke, Hani Goodarzi, Patrick D. Hsu, and Brian L. Hie. Genome modeling and design across all domains of life with Evo 2, February 2025.

9. Antonio Pedro Camargo, Simon Roux, Frederik Schulz, Michal Babinski, Yan Xu, Bin Hu, Patrick S. G. Chain, Stephen Nayfach, and Nikos C. Kyrpides. Identification of mobile genetic elements with geNomad. Nature Biotechnology, 42(8):1303–1312, August 2024.

10. Carlos Canchaya, Ghislain Fournous, and Harald Brüssow. The impact of prophages on bacterial chromosomes. Molecular Microbiology, 53(1):9–18, 2004.

11. Sherwood Casjens. Prophages and bacterial genomics: what have we learned so far? Molecular Microbiology, 49(2):277–300, July 2003.

12. Wenduo Cheng, Zhenqiao Song, Yang Zhang, Shike Wang, Danqing Wang, Muyu Yang, Lei Li, and Jian Ma. DNALONGBENCH: a benchmark suite for long-range DNA prediction tasks. Nature Communications, 16(1):10108, November 2025.

13. Julien Colin, Lore Goetschalckx, Thomas Fel, Victor Boutin, Jay Gopal, Thomas Serre, and Nuria Oliver. Local vs distributed representations: What is the right basis for interpretability?, November 2024. arXiv:2411.03993 [cs].

14. Ryan Cook, Nathan Brown, Tamsin Redgwell, Branko Rihtman, Megan Barnes, Martha Clokie, Dov J. Stekel, Jon Hobman, Michael A. Jones, and Andrew Millard. INfrastructure for a PHAge REference Database: Identification of Large-Scale Biases in the Current Collection of Cultured Phage Genomes. PHAGE, 2(4):214–223, December 2021.

15. Hoagy Cunningham, Aidan Ewart, Logan Riggs, Robert Huben, and Lee Sharkey. Sparse Autoencoders Find Highly Interpretable Features in Language Models, October 2023. arXiv:2309.08600 [cs].

16. Hugo Dalla-Torre, Liam Gonzalez, Javier Mendoza-Revilla, Nicolas Lopez Carranza, Adam Henryk Grzywaczewski, Francesco Oteri, Christian Dallago, Evan Trop, BernardoP. de Almeida, Hassan Sirelkhatim, Guillaume Richard, Marcin Skwark, Karim Beguir, Marie Lopez, and Thomas Pierrot. The Nucleotide Transformer: building and evaluating robust foundation models for human genomics, October 2024. bioRxiv: 10.1101/2023.01.11.523679.

17. Moïra B. Dion, Frank Oechslin, and Sylvain Moineau. Phage diversity, genomics and phylogeny. Nature Reviews Microbiology, 18(3):125–138, March 2020. Number: 3.

18. Veniamin Fishman, Yuri Kuratov, Aleksei Shmelev, Maxim Petrov, Dmitry Penzar, Denis Shepelin, Nikolay Chekanov, Olga Kardymon, and Mikhail Burtsev. GENA-LM: a family of open-source foundational DNA language models for long sequences. Nucleic Acids Research, 53(2):gkae1310, January 2025.

19. Hongyan Gao, Bowen Li, Zihan Guo, Lei Zheng, Junnan Chen, and Guanxiang Liang. Highly accurate prophage island detection with PIDE. Genome Biology, 26:254, August 2025.

20. Christian H Gauthier, Lawrence Abad, Ananya K Venbakkam, Julia Malnak, Daniel A Russell, and Graham F Hatfull. DEPhT: a novel approach for efficient prophage discovery and precise extraction. Nucleic Acids Research, 50(13):e75–e75, July 2022.

21. Susanna R. Grigson, George Bouras, Bas E. Dutilh, Robert D. Olson, and Robert A. Edwards. Computational function prediction of bacteria and phage proteins. Microbiology and molecular biology reviews: MMBR, 89(3):e0002225, September 2025.

22. Jiarong Guo, Ben Bolduc, Ahmed A. Zayed, Arvind Varsani, Guillermo Dominguez-Huerta, Tom O. Delmont, Akbar Adjie Pratama, M. Consuelo Gazitúa, Dean Vik, Matthew B. Sullivan, and Simon Roux. VirSorter2: a multi-classifier, expert-guided approach to detect diverse DNA and RNA viruses. Microbiome, 9(1):37, February 2021.

23. Wes Gurnee, Neel Nanda, Matthew Pauly, Katherine Harvey, Dmitrii Troitskii, and Dimitris Bertsimas. Finding Neurons in a Haystack: Case Studies with Sparse Probing, June 2023. arXiv:2305.01610 [cs].

24. Iain D. Hay and Trevor Lithgow. Filamentous phages: masters of a microbial sharing economy. EMBO Reports, 20(6), June 2019.

25. Dan Hendrycks, Collin Burns, Steven Basart, Andy Zou, Mantas Mazeika, Dawn Song, and Jacob Steinhardt. Measuring Massive Multitask Language Understanding, January 2021. arXiv:2009.03300 [cs].

26. Yanrong Ji, Zhihan Zhou, Han Liu, and Ramana V Davuluri. DNABERT: pre-trained bidirectional encoder representations from transformers model for DNA-language in genome. Bioinformatics, 37:2112–2120, August 2021.

27. Vanessa Isabell Jurtz, Julia Villarroel, Ole Lund, Mette Voldby Larsen, and Morten Nielsen. MetaPhinder—Identifying Bacteriophage Sequences in Metagenomic Data Sets. PLOS ONE, 11(9):e0163111, September 2016.

28. Kristopher Kieft, Zhichao Zhou, and Karthik Anantharaman. VIBRANT: automated recovery, annotation and curation of microbial viruses, and evaluation of viral community function from genomic sequences. Microbiome, 8(1):90, June 2020.

29. James C. Kosmopoulos and Karthik Anantharaman. Viral Dark Matter: Illuminating Protein Function, Ecology, and Biotechnological Promises. Biochemistry, 64(24):4609–4627, December 2025.

30. Qiuyi Li, Wei Wu, Yiheng Zhu, Fuli Feng, Jieping Ye, and Zheng Wang. Generanno: A Genomic Foundation Model for Metagenomic Annotation, August 2025. ISSN: 2692-8205 Pages: 2025.06.04.656517 Section: New Results.

31. Balázs Ligeti, István Szepesi-Nagy, Balázs Bodnár, Nóra Ligeti-Nagy, and József Juhász. ProkBERT family: genomic language models for microbiome applications. Frontiers in Microbiology, 14:1331233, January 2024.

32. Gita Mahmoudabadi and Rob Phillips. A comprehensive and quantitative exploration of thousands of viral genomes. eLife, 7:e31955, April 2018.

33. Frederikke Isa Marin, Felix Teufel, Marc Horlacher, Dennis Madsen, Dennis Pultz, Ole Winther, and Wouter Boomsma. BEND: Benchmarking DNA Language Models on biologically meaningful tasks, April 2024. arXiv:2311.12570 [q-bio].

34. Aashiq Muhamed, Mona Diab, and Virginia Smith. Decoding Dark Matter: Specialized Sparse Autoencoders for Interpreting Rare Concepts in Foundation Models, November 2024. arXiv:2411.00743 [cs].

35. Hugo Oliveira, Rita Domingues, Benjamin Evans, J. Mark Sutton, Evelien M. Adriaenssens, and Dann Turner. Genomic diversity of bacteriophages infecting the genus Acinetobacter. Viruses, 14(2):181, January 2022.

36. Donovan H Parks, Maria Chuvochina, Christian Rinke, Aaron J Mussig, Pierre-Alain Chaumeil, and Philip Hugenholtz. GTDB: an ongoing census of bacterial and archaeal diversity through a phylogenetically consistent, rank normalized and complete genome-based taxonomy. Nucleic Acids Research, 50(D1):D785–D794, January 2022.

37. Donovan H. Parks, Michael Imelfort, Connor T. Skennerton, Philip Hugenholtz, and Gene W. Tyson. CheckM: assessing the quality of microbial genomes recovered from isolates, single cells, and metagenomes. Genome Research, 25(7):1043–1055, July 2015.

38. Jie Ren, Nathan A. Ahlgren, Yang Young Lu, Jed A. Fuhrman, and Fengzhu Sun. VirFinder: a novel k-mer based tool for identifying viral sequences from assembled metagenomic data. Microbiome, 5(1):69, July 2017.

39. Jie Ren, Kai Song, Chao Deng, Nathan A. Ahlgren, Jed A. Fuhrman, Yi Li, Xiaohui Xie, Ryan Poplin, and Fengzhu Sun. Identifying viruses from metagenomic data using deep learning. Quantitative Biology (Beijing, China), 8(1):64–77, March 2020.

40. Reza Rezaei Javan, Elisa Ramos-Sevillano, Asma Akter, Jeremy Brown, and Angela B. Brueggemann. Prophages and satellite prophages are widespread in Streptococcus and may play a role in pneumococcal pathogenesis. Nature Communications, 10(1):4852, October 2019.

41. Yair Schiff, Chia-Hsiang Kao, Aaron Gokaslan, Tri Dao, Albert Gu, and Volodymyr Kuleshov. Caduceus: Bi-Directional Equivariant Long-Range DNA Sequence Modeling. In Proceedings of Machine Learning Research, volume 235, pages 43632–43648, 2024.

42. Bin Shao and Jiawei Yan. A long-context language model for deciphering and generating bacteriophage genomes. Nature Communications, 15(1):9392, October 2024.

43. Kimmo Sirén, Andrew Millard, Bent Petersen, M Thomas P Gilbert, Martha R J Clokie, and Thomas Sicheritz-Pontén. Rapid discovery of novel prophages using biological feature engineering and machine learning. NAR Genomics and Bioinformatics, 3(1):lqaa109, March 2021.

44. Elizaveta V Starikova, Polina O Tikhonova, Nikita A Prianichnikov, Chris M Rands, Evgeny M Zdobnov, Elena N Ilina, and Vadim M Govorun. Phigaro: high-throughput prophage sequence annotation. Bioinformatics, 36(12):3882–3884, June 2020.

45. Ziqi Tang, Nirali Somia, Yiyang Yu, and Peter K. Koo. Evaluating the representational power of pre-trained DNA language models for regulatory genomics. Genome Biology, 26(1):203, July 2025.

46. Kirill Vishniakov, Karthik Viswanathan, Aleksandr Medvedev, Praveen K. Kanithi, Marco AF Pimentel, Ronnie Rajan, and Shadab Khan. Genomic Foundationless Models: Pretraining Does Not Promise Performance, December 2024. Pages: 2024.12.18.628606 Section: New Results.

47. Kirill Vishniakov, Karthik Viswanathan, Aleksandr Medvedev, Praveenkumar Kanithi, Marco AF Pimentel, Ronnie Rajan, and Shadab Khan. Genomic Foundationless Models: Pretraining Does Not Promise Performance, June 2025. Pages: 2024.12.18.628606 Section: New Results.

48. Alex Wang, Yada Pruksachatkun, Nikita Nangia, Amanpreet Singh, Julian Michael, Felix Hill, Omer Levy, and Samuel R. Bowman. SuperGLUE: A Stickier Benchmark for General-Purpose Language Understanding Systems, February 2020. arXiv:1905.00537 [cs].

49. Alex Wang, Amanpreet Singh, Julian Michael, Felix Hill, Omer Levy, and Samuel R. Bowman. GLUE: A Multi-Task Benchmark and Analysis Platform for Natural Language Understanding, February 2019. arXiv:1804.07461 [cs].

50. Manzil Zaheer, Guru Guruganesh, Avinava Dubey, Joshua Ainslie, Chris Alberti, Santiago Ontanon, Philip Pham, Anirudh Ravula, Qifan Wang, Li Yang, and Amr Ahmed. Big bird: transformers for longer sequences. In Proceedings of the 34th International Conference on Neural Information Processing Systems, NIPS ‘20, Red Hook, NY, USA, 2020. Curran Associates Inc.

51. You Zhou, Yongjie Liang, Karlene H. Lynch, Jonathan J. Dennis, and David S. Wishart. PHAST: A fast phage search tool. Nucleic Acids Research, 39(Web Server issue):W347–W352, July 2011.

52. Zhihan Zhou, Yanrong Ji, Weijian Li, Pratik Dutta, Ramana Davuluri, and Han Liu. DNABERT-2: Efficient Foundation Model and Benchmark For Multi-Species Genome, March 2024. arXiv:2306.15006 [q-bio].

53. Andrzej Zielezinski, Adam Gudyś, Jakub Barylski, Krzysztof Siminski, Piotr Rozwalak, Bas E. Dutilh, and Sebastian Deorowicz. Ultrafast and accurate sequence alignment and clustering of viral genomes. Nature Methods, 22(6):1191– 1194, June 2025.

