## Supplementary material for "LAMBDA: A Prophage Detection Benchmark for Genomic Language Models": LAMBDA v1 Supplementary Figures

### Supplementary Table Legend

**S1: Genome-Wide Prophage Detection Evaluation Set**

**S2: Manual Review of Novel Candidate Prophage Regions**

**S3: PHROG Functional Category Sensitivity by Model**

**S4: Optimal Hyperparameters for Prophage Signal Extraction**

### Supplementary Figures Legend

**Figure S1: Hyperparameter Grid-Search for Prophage Signal Extraction Algorithm**

**Figure S2: Pretrained Embedding Performance Compared to Random for 2k, 4k, 8k Datasets**

**Figure S3: Case study of clean genome-wide prophage detection in *Mycobacteroides abscessus subsp. abscessus strain GD17 (GCA_017183835.1) with average MCC score of 0.945***

**Figure S4: Case study of genome-wide prophage detection in *Escherichia coli* K-12 (NC_000913) with an average gLM MCC score of 0.198.**

**Figure S5: Case study of genome-wide prophage detection in *Pseudomonas aeruginosa MTB-1* (GCA_000504045.1) with an average gLM MCC score of 0.696.**

**Figure S6: Detailed view of Candidate Prophage #19 (663,168– 704,000 bp) in *Pseudomonas aeruginosa MTB-1* (GCA_000504045.1) with PHROG functional annotations.**

**Figure S7: Model Performance Stratified by Bacterial Genome Taxonomy (Phylum)**

**Figure S8: Model Performance Stratified by Bacterial Genome Taxonomy (Class)**

**Figure S9: Model Performance Stratified by Bacterial Genome Taxonomy (Order)**

**Figure S10: Model Performance Stratified by Bacterial Genome Taxonomy (Genus)**

**Figure S11: Model Performance Stratified by Viral Lineage**

**Figure S12: Model Performance compared to MASH distance to Nearest INPHARED Phage**

**Figure S13: Per-model Detection Rate for Known vs. Novel Prophages**

**Figure S14: Detection support for reference prophages compared to candidate regions**

**Figure S15: Prophage detectability rises with representation in the phage training set**

**Figure S16: Case study of genome-wide prophage detection in *Mycobacteroides abscessus* subsp. *abscessus* strain GD43A (EVO2 recall = 1.00; EVO2+SAE recall = 0.98)**

**Figure S17: Case study of genome-wide prophage detection in *Escherichia coli* O157:H7 str. Sakai (EVO2 recall = 0.82; EVO2+SAE recall = 0.38)**

**Figure S18: Case study of genome-wide prophage detection in *Wolbachia* endosymbiont of *Drosophila melanogaster* (EVO2 recall = 0.47; EVO2+SAE recall = 0.00)**

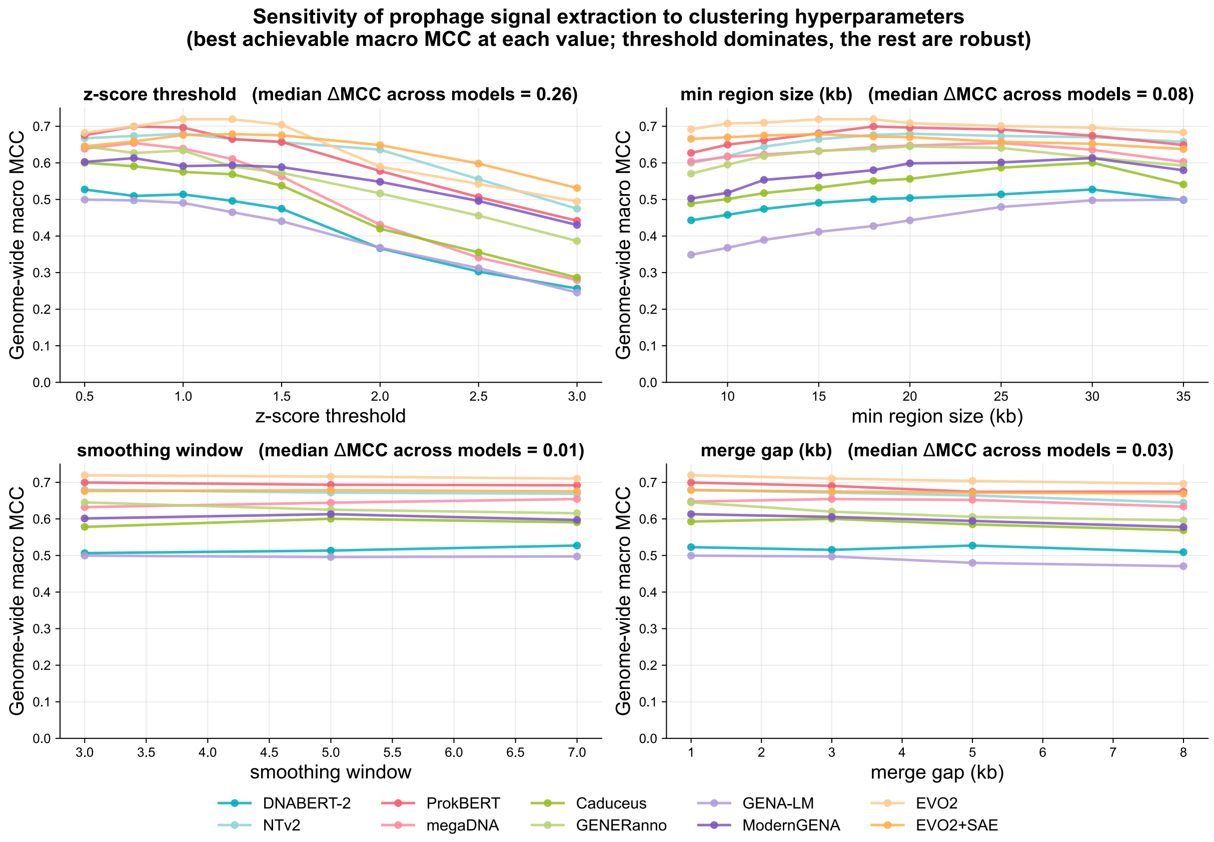


**Figure S1: Hyperparameter Grid-Search for Prophage Signal Extraction Algorithm.** Predicted regions were filtered by score threshold (y-axis), defined as the minimum average per-position prophage probability, and minimum region size (x-axis). Each cell reports the resulting genome-wide MCC for that parameter combination, with the best MCC per model shown in bold. Optimal filtering parameters for each model were used to report the genome wide phage prediction results.

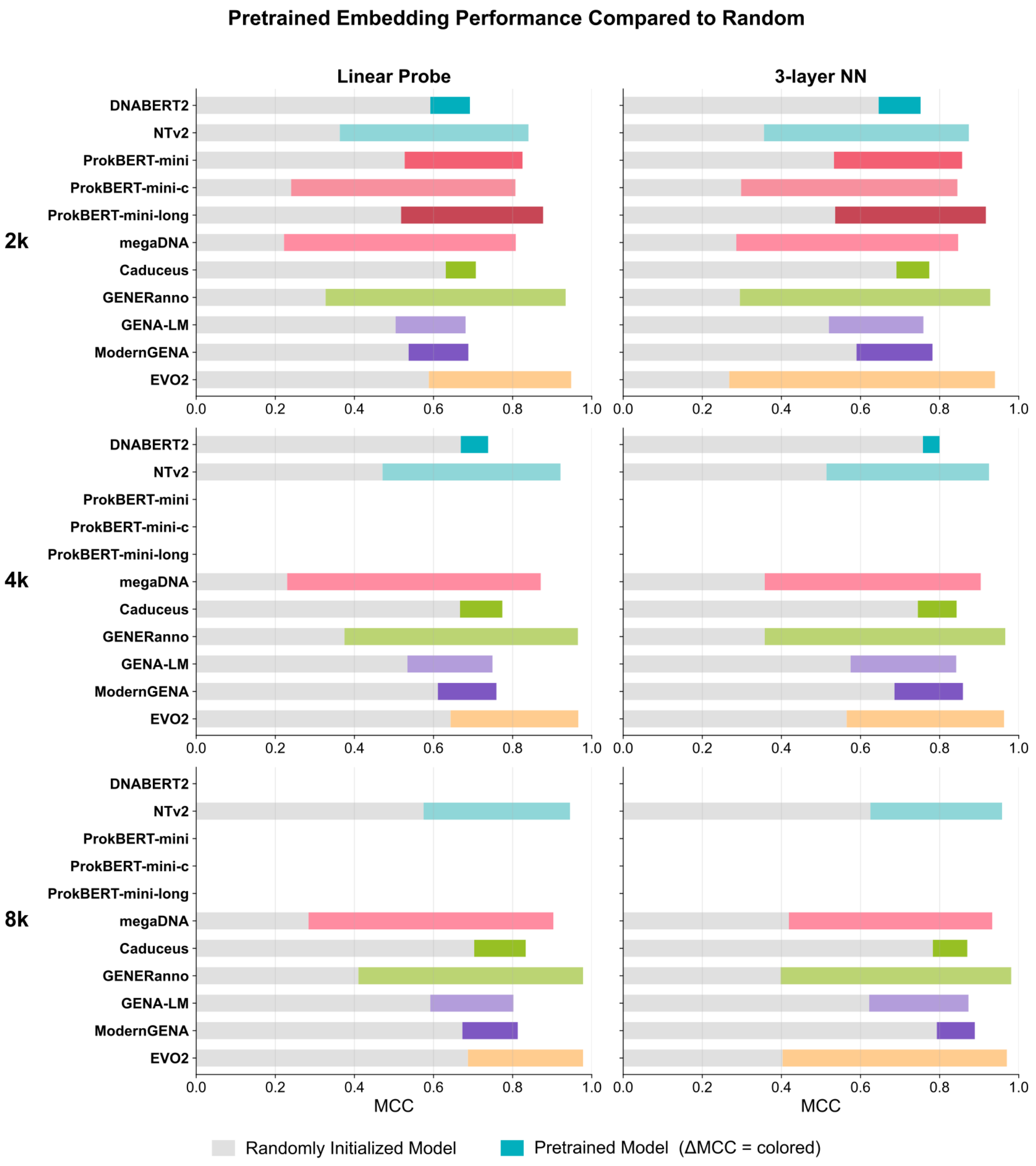


**Figure S2: Pretrained Embedding Performance Compared to Random for 2k, 4k, 8k Datasets.** Pretrained embedding strength is defined as the difference in MCC between pretrained embeddings and embeddings from the same model with randomly initialized weights (ΔMCC), where larger ΔMCC values indicate a stronger contribution from pretraining. Models were only evaluated on sequences that fit in their context window.

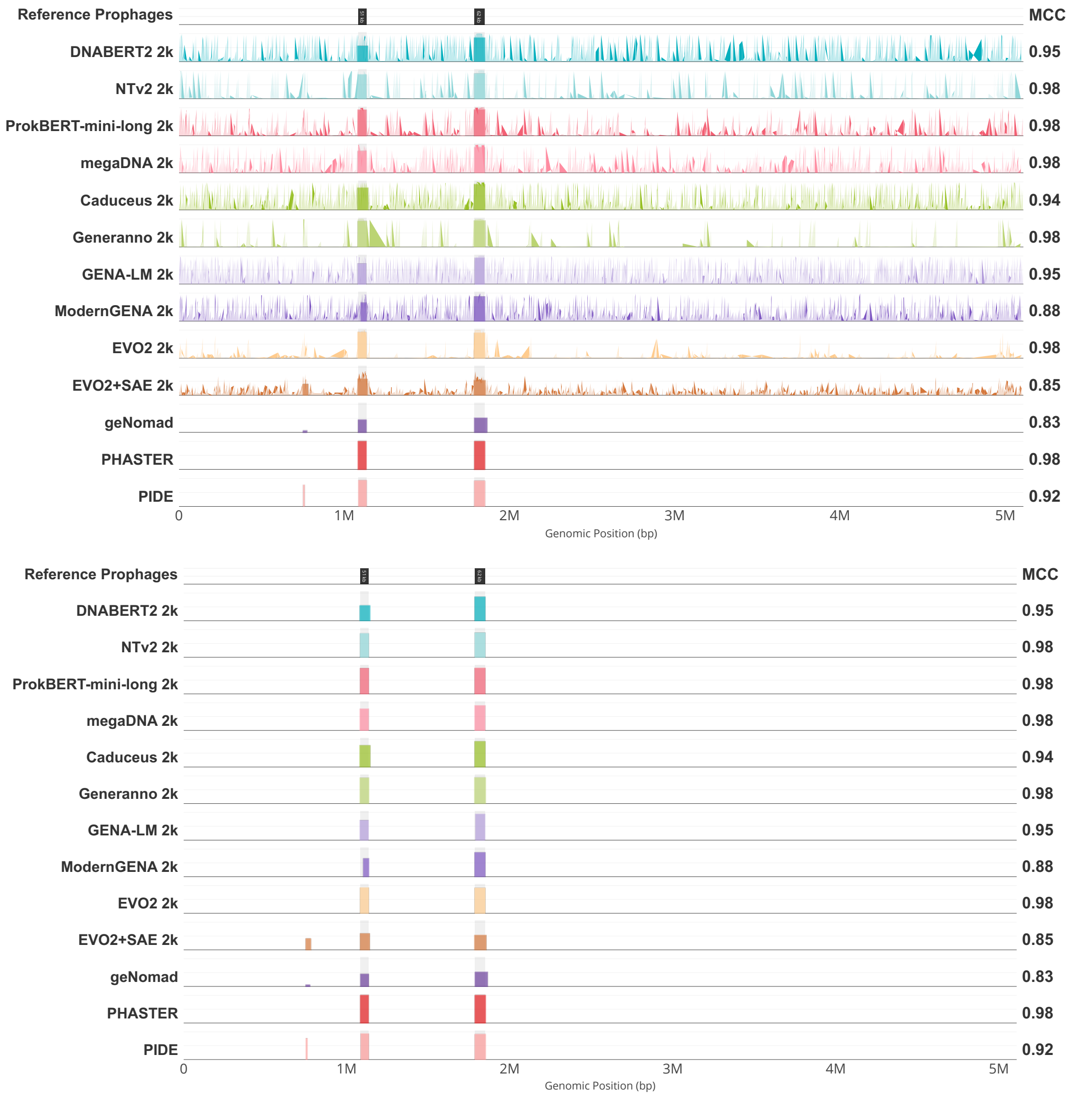


**Figure S3: Case study of clean genome-wide prophage detection in *Mycobacteroides abscessus subsp. abscessus strain GD17 (GCA_017183835.1) with average MCC score of 0.945.*** Raw embedding-derived signal (top), and predicted regions (bottom). In this example, strong, localized peaks in the raw model signal align closely with annotated prophage loci, producing sharp, well-delimited predictions with minimal background activation across the rest of the genome. Similar interactive visualizations are available for all LAMBDA test genomes at https://leannmlindsey.github.io/lambda-benchmark/.

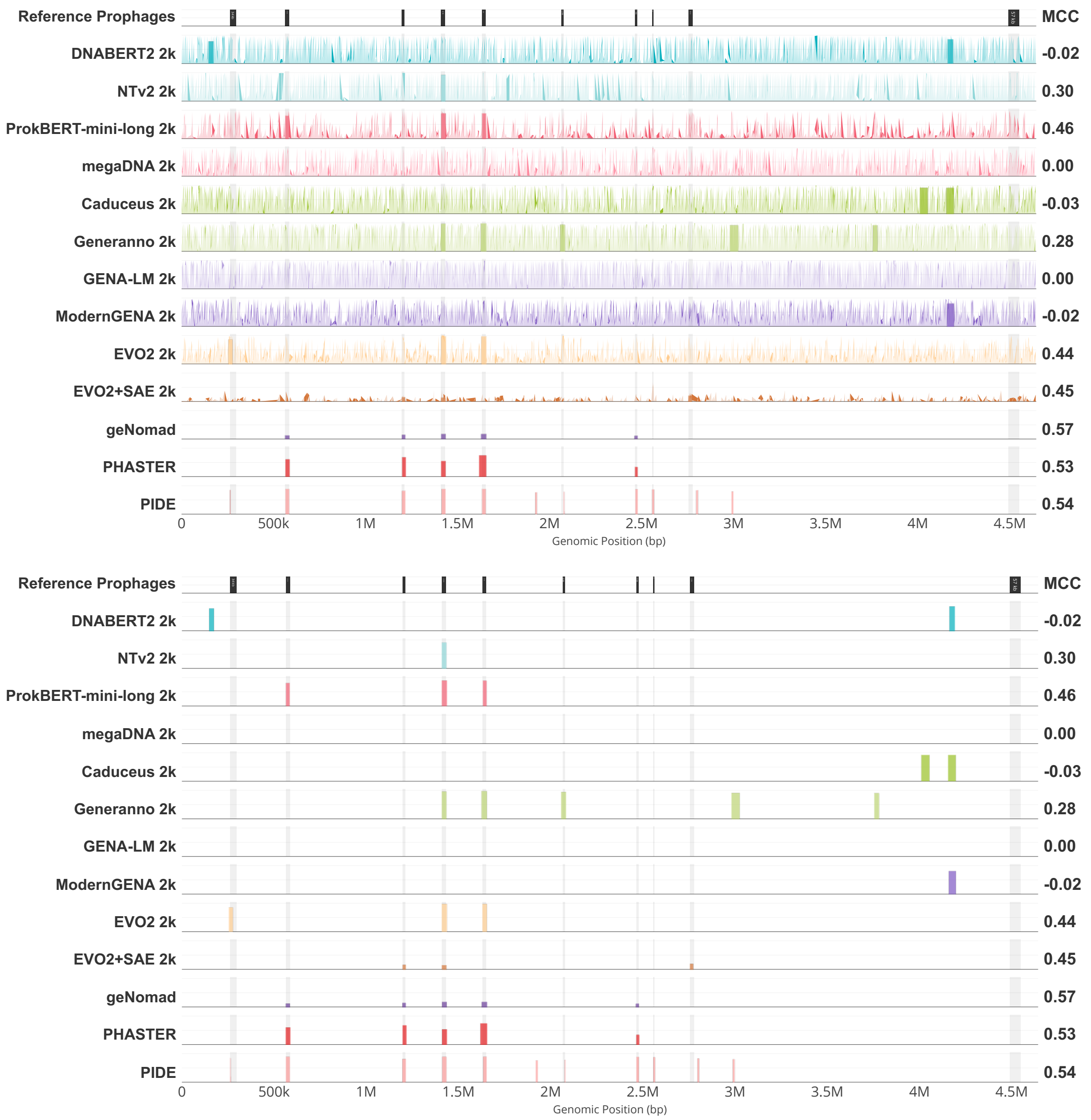


**Figure S4: Case study of genome-wide prophage detection in *Escherichia coli* K-12 (NC_000913) with an average gLM MCC score of 0.198.** Raw embedding-derived signal (top), and predicted regions (bottom). In contrast to the *Mycobacterioides abscessus* example (Figure S3), the prophage signal is low, combined with a high false positive baseline, resulting in poor detection by most models. Similar interactive visualizations are available for all LAMBDA test genomes at https://leannmlindsey.github.io/lambda-benchmark/.

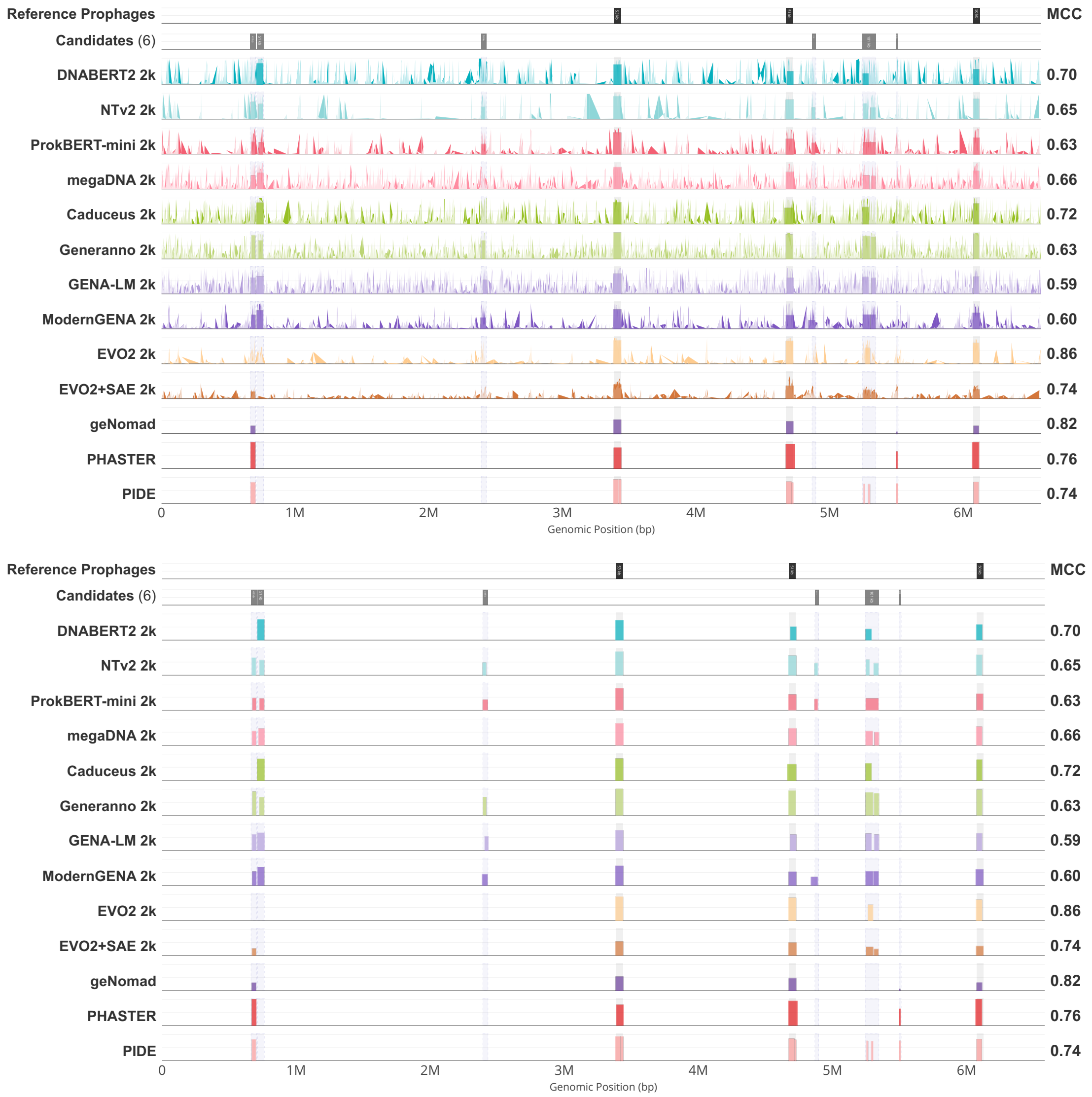


**Figure S5: Case study of genome-wide prophage detection in *Pseudomonas aeruginosa MTB-1* (GCA_000504045.1) with an average gLM MCC score of 0.696.** Raw embedding-derived signal (top), and predicted regions (bottom). Six candidate prophage regions are identified that highlight cross-method agreement. Similar interactive visualizations are available for all LAMBDA test genomes at https://leannmlindsey.github.io/lambda-benchmark/.

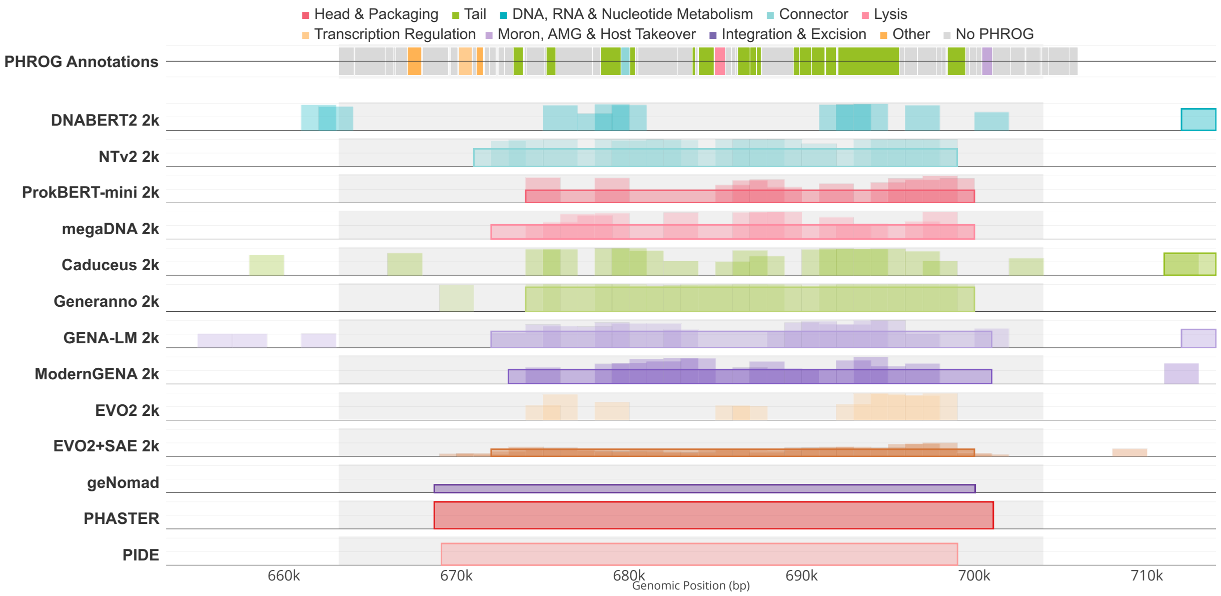


**Figure S6: Detailed view of Candidate Prophage #19 (663,168– 704,000 bp) in *Pseudomonas aeruginosa MTB-1* (GCA_000504045.1) with PHROG functional annotations.** Zoomed-in visualization of prophage region 1 (±10 kb padding) showing PHROG functional categories across genes (top track) alongside genomic language model predictions and traditional tool calls. It is supported by 14 detection methods and classified as a likely phage, with strong structural evidence with 51% of its genes encoding phage structural proteins (head, tail, connector, and lysis). Similar interactive visualizations are available for all prophage regions in the benchmark at https://leannmlindsey.github.io/lambda-benchmark/

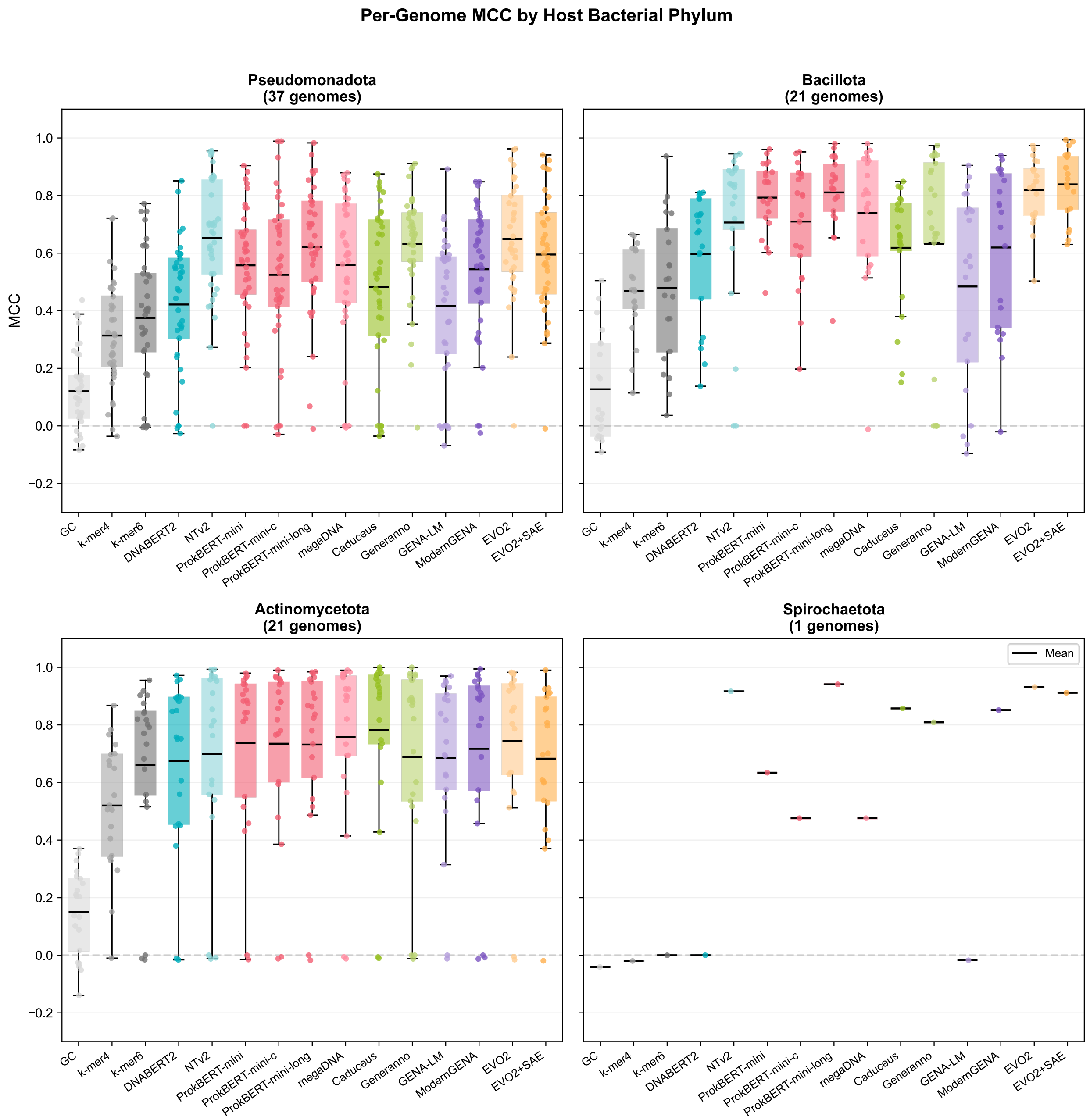


**Figure S7: Model Performance Stratified by Bacterial Genome Taxonomy (Phylum).** Per-genome MCC of prophage detection by host bacterial phylum. Each panel shows one phylum, with boxplots for each model (2k input). Points represent individual genomes. Black lines indicate means.

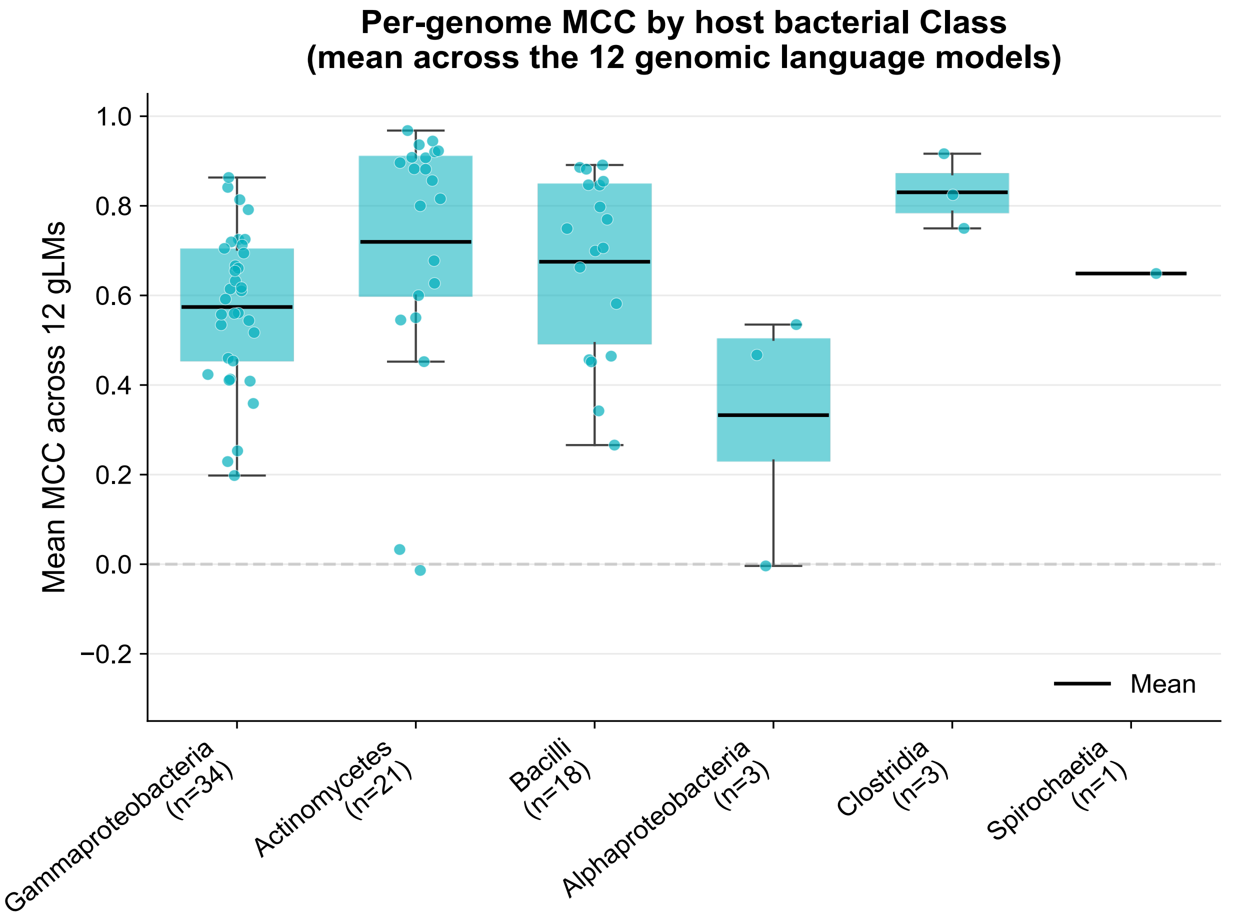


**Figure S8: Model Performance Stratified by Bacterial Genome Taxonomy (Class).** Per-genome MCC of prophage detection by host bacterial class. Each panel shows one phylum, with boxplots for each model (2k input). Points represent individual genomes. Black lines indicate means.

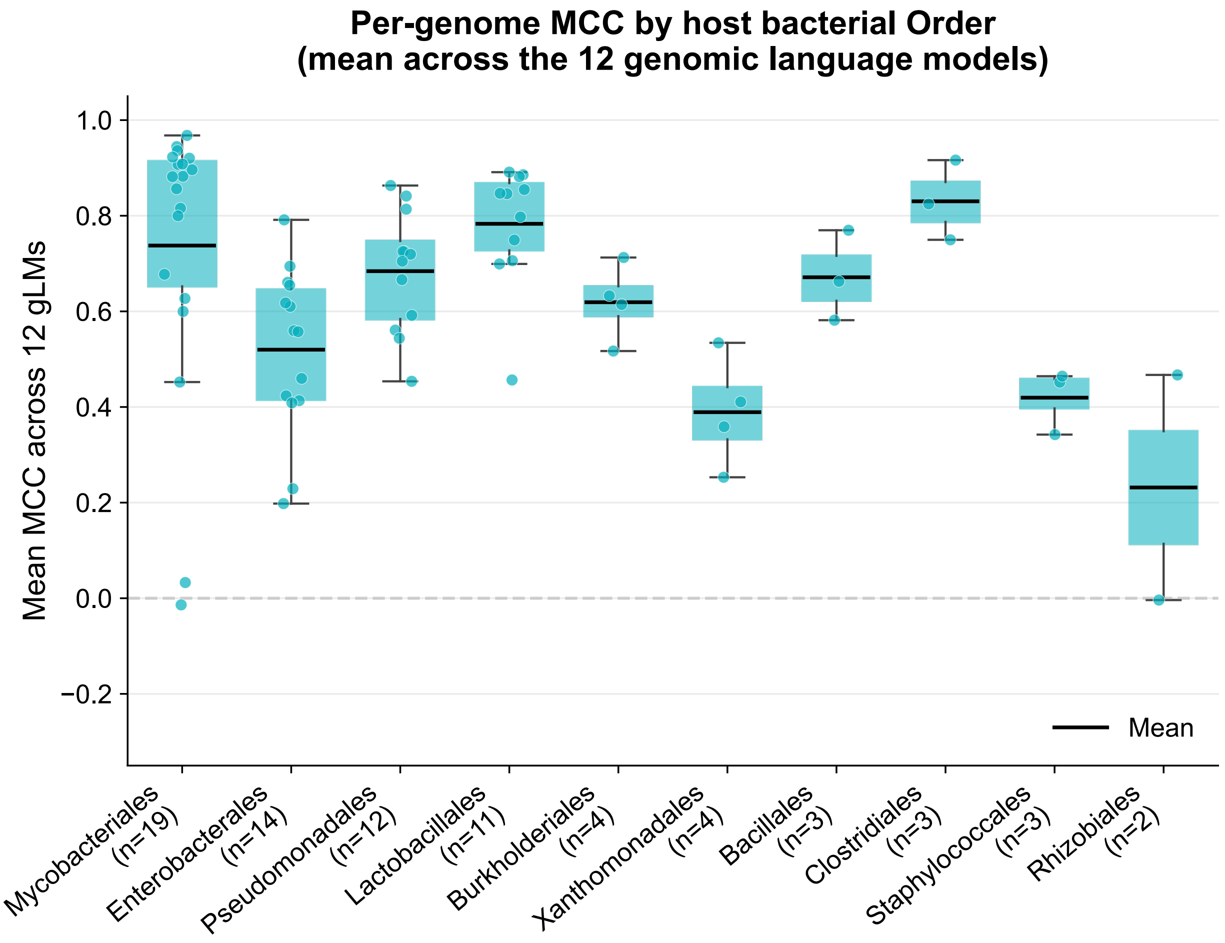


**Figure S9: Model Performance Stratified by Bacterial Genome Taxonomy (Order).** Per-genome MCC of prophage detection by host bacterial order. Groups are sorted by number of genomes (descending). Singletons are included at the end.

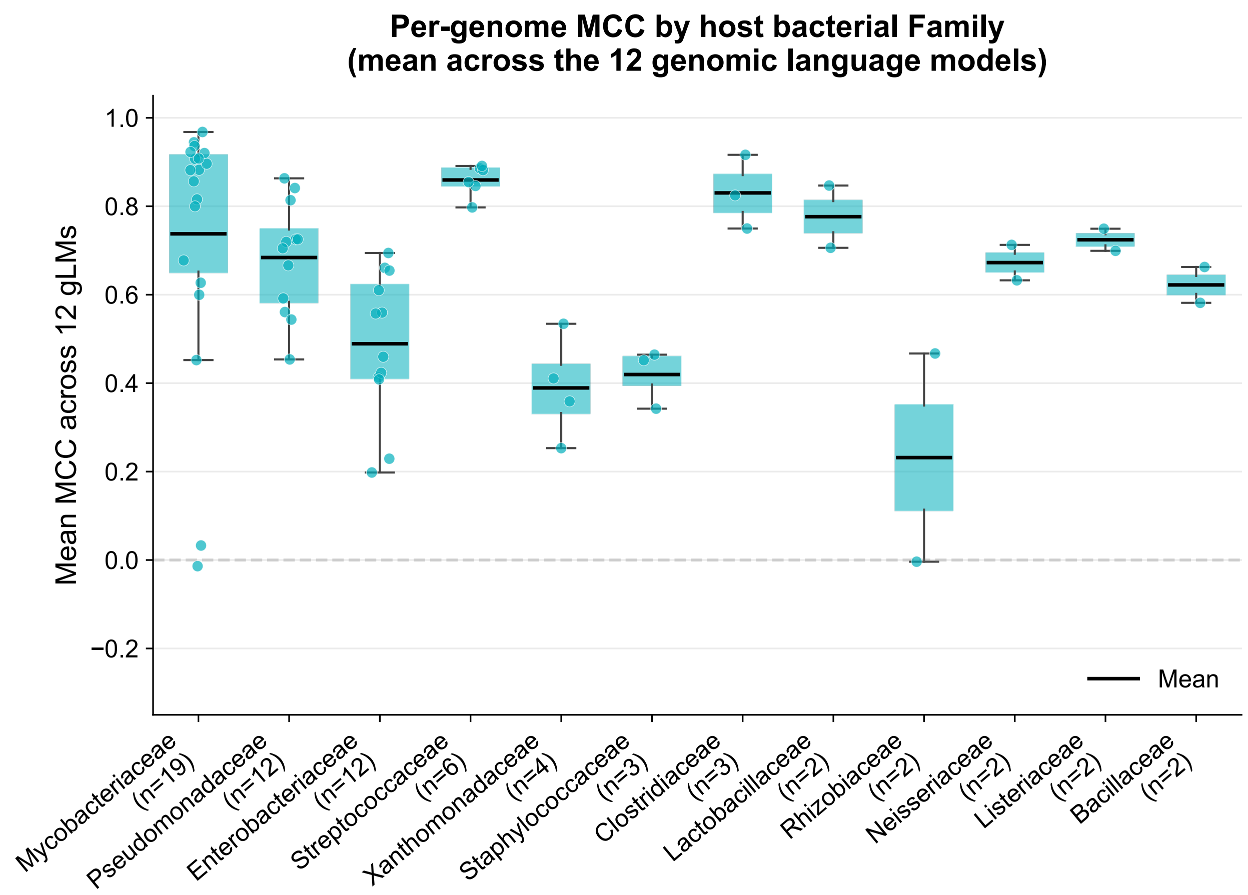


**Figure S10: Model Performance Stratified by Bacterial Genome Taxonomy (Genus).** Per-genome MCC of prophage detection by host bacterial genus (genera with n≥2 genomes). Groups are sorted by number of genomes (descending).

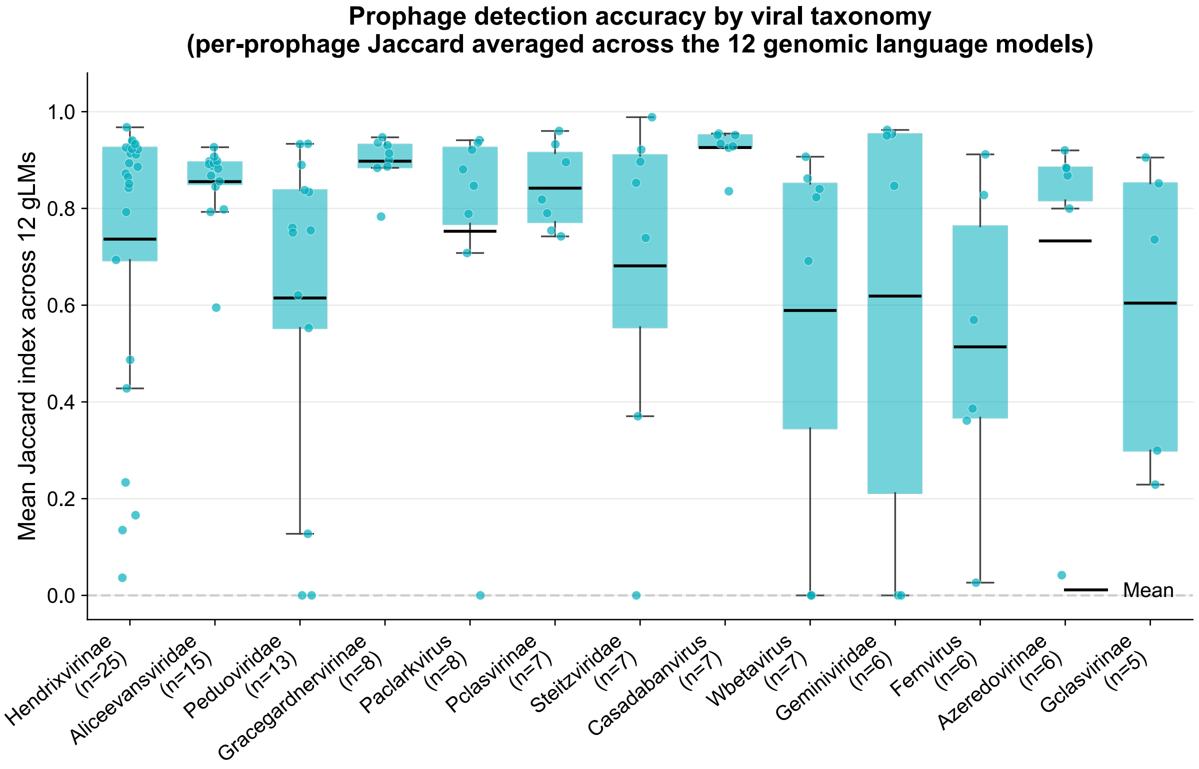


**Figure S11: Model Performance Stratified by Viral Lineage.** Per-prophage Jaccard index by viral lineage (lineages with n≥5 prophages). Lineage classifications were assigned by Pharokka. Higher Jaccard values indicate better spatial overlap between predicted and ground-truth prophage boundaries.

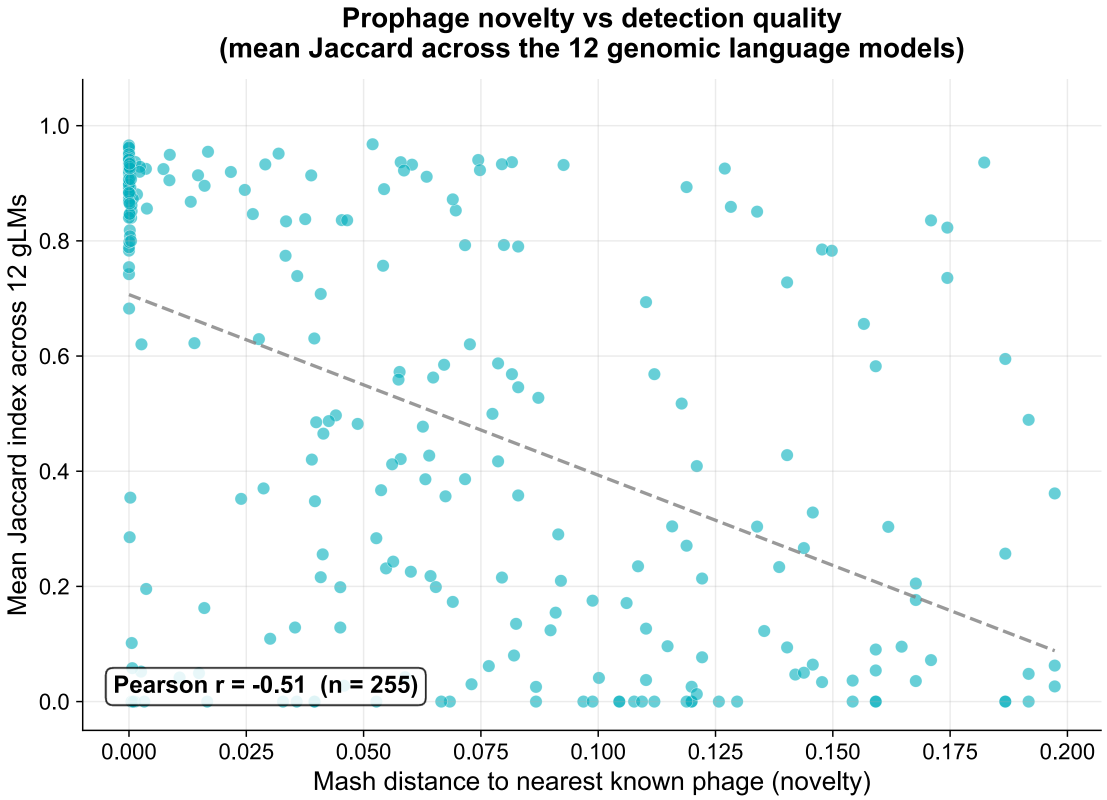


**Figure S12: Model Performance compared to MASH distance to Nearest INPHARED Phage.** Each panel shows the per-prophage Jaccard Index as a function of Mash distance to the nearest known phage in the INPHARED database, as determined by Pharokka. Individual prophages are shown as points, with lines connecting binned means to highlight trends. Pearson correlation coefficients (r) are reported for each model. All models with sufficient predictions show a negative correlation, indicating that prophages more distant from characterized phages are detected less accurately.

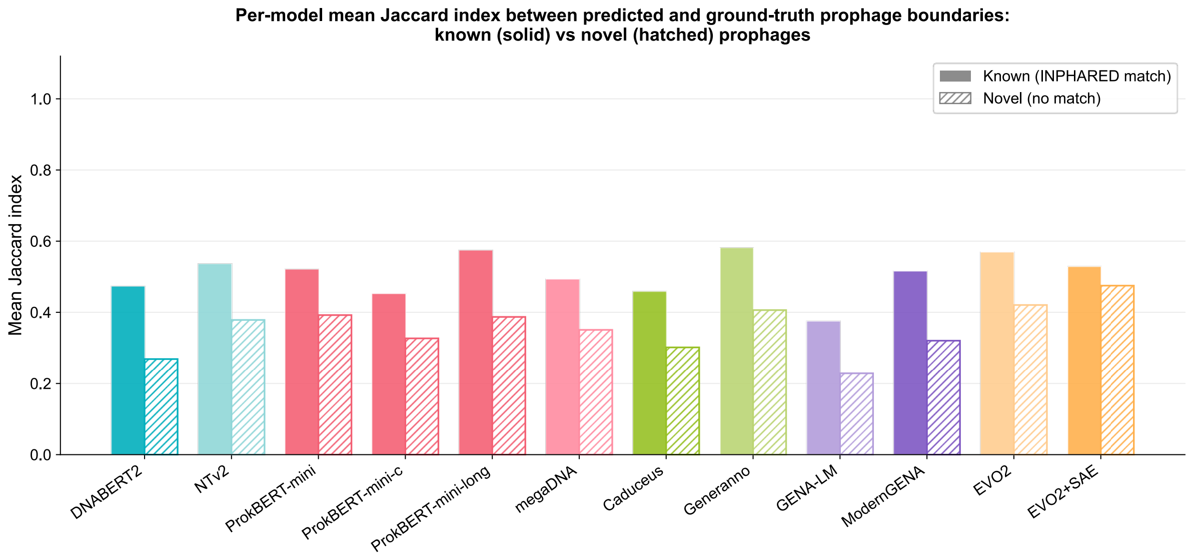


**Figure S13: Per-model Mean Jaccard Index for Known vs. Novel Prophages.** Per-model detection rate for known vs novel prophages. Mean Jaccard Index between predicted and ground-truth prophage boundaries for prophages with a match in the INPHARED database (solid bars "Known") versus those with no match (hatched bars, "Novel"), Novelty was determined using Mash distance threshold of greater than 0.2 as computed by Pharokka. All models show reduced detection quality on novel prophages, with Generanno showing the smallest performance gap (0.36 vs 0.33)

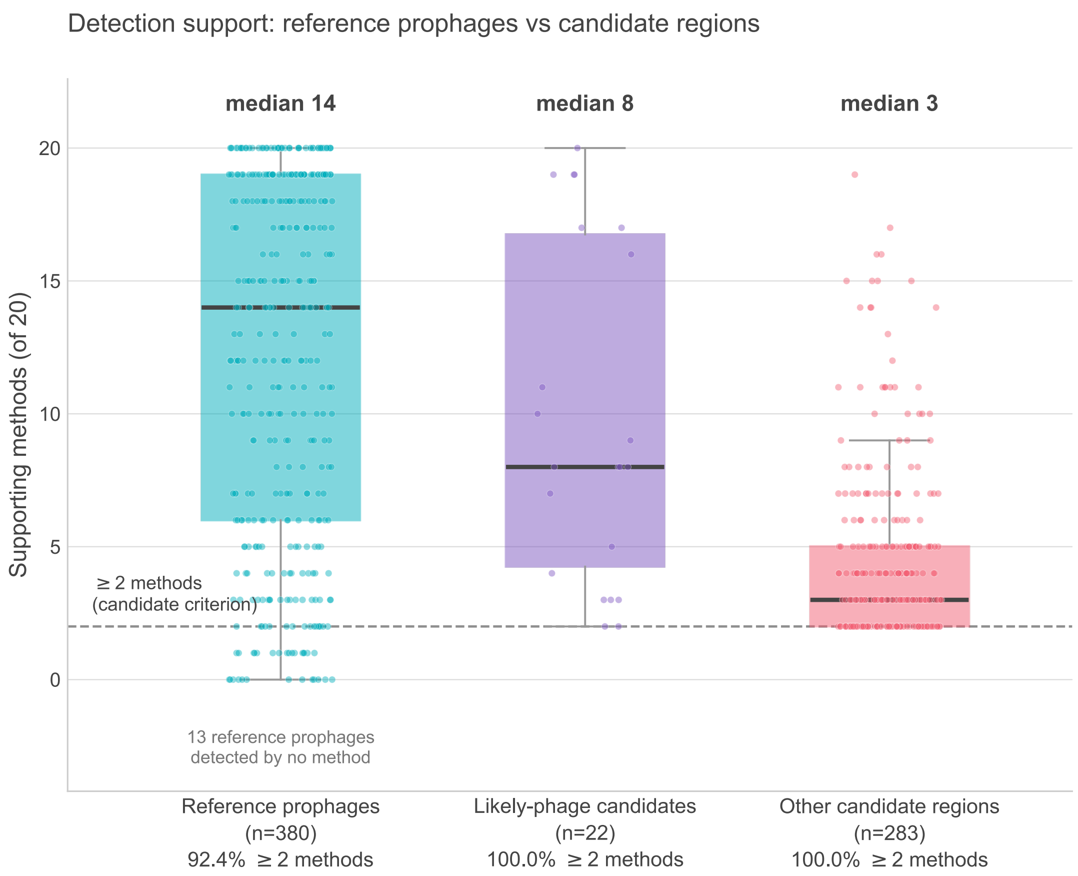


**Figure S14. Detection support for reference prophages compared to candidate regions.** Support is the number of the 20 detection methods (12 gLMs, 8 dedicated tools) recovering a region in the genome-wide scan. Reference prophages had a median support of 14, with 351 of 380 (92.4%) meeting the ≥2-method criterion used to define candidates (dashed line); 13 were detected by no method (Supplementary Material Figure S15). Likely-phage candidates had a median of 8, other candidate regions a median of 3. Boxes show the interquartile range and median; points are individual regions.

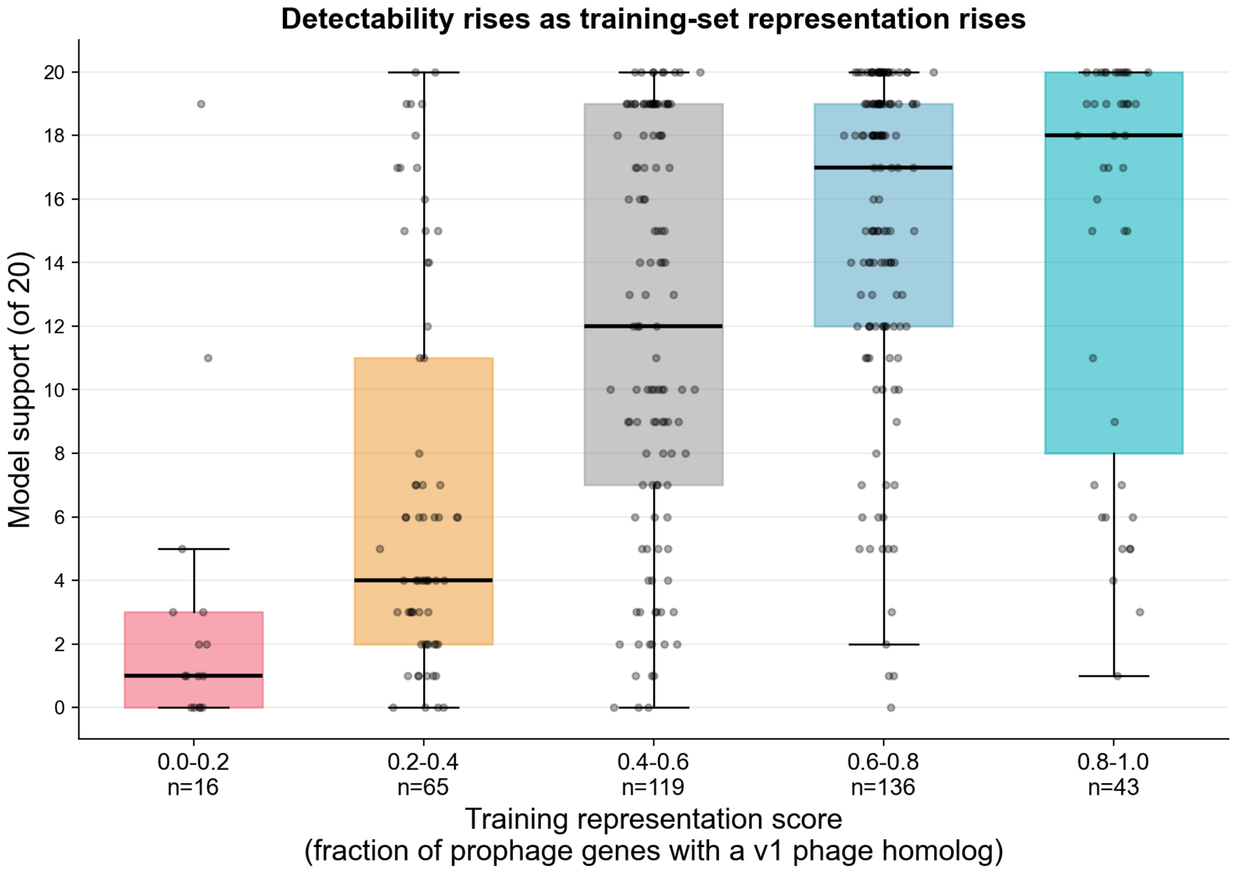


**Figure S15. Prophage detectability rises with representation in the phage training set.** The 380 reference prophages were grouped into five bins by training-representation score, defined as the fraction of a prophage's predicted genes with a protein-level homolog (BLASTX, $E < 10^{-5}$) among the LAMBDA phage training sequences. Each box shows the distribution of model support, the number of the 20 models and tools that detected that prophage in the genome-wide scan. Median support rises monotonically across bins, from 1 of 20 among the least-represented prophages to 18 of 20 among the most-represented (Spearman rho = 0.48, $p = 9.5 \times 10^{-24}$).

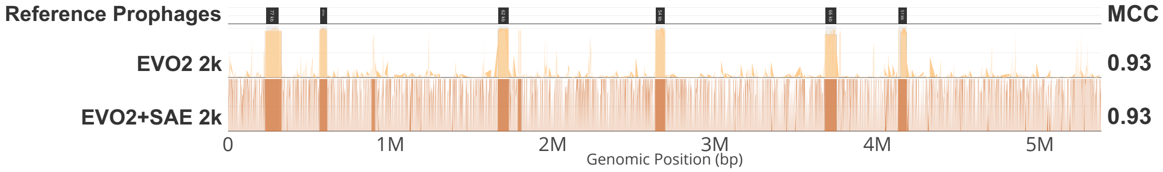


**Figure S16: Case study of genome-wide prophage detection in *Mycobacteroides abscessus* subsp. *abscessus* strain GD43A (EVO2 recall = 1.00; EVO2+SAE recall = 0.98).** Genome-wide tracks comparing raw EVO2 embedding-derived signal (yellow) with activations from the sparse autoencoder applied to EVO2 (orange). Published ground truth prophage regions are shown in black at the top of the panel. EVO2+SAE achieves recall nearly equivalent to EVO2 but with slightly lower precision (0.75 vs. 0.92). This example illustrates a case where the SAE feature largely captures the prophage signal.

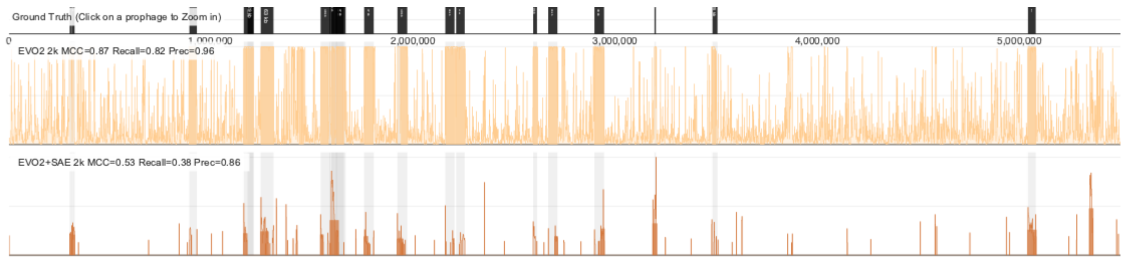


**Figure S17: Case study of genome-wide prophage detection in *Escherichia coli* O157:H7 str. Sakai (EVO2 recall = 0.82; EVO2+SAE recall = 0.38).** Genome-wide tracks comparing raw EVO2 embedding-derived signal (yellow) with sparse autoencoder activations (orange), with published ground truth prophage regions shown in black. The SAE signal is substantially sparser, with reduced recall reflecting incomplete activation across full prophage regions and limited separation between true signal and false positives. This example illustrates a case where the SAE feature only partially captures the prophage signal.

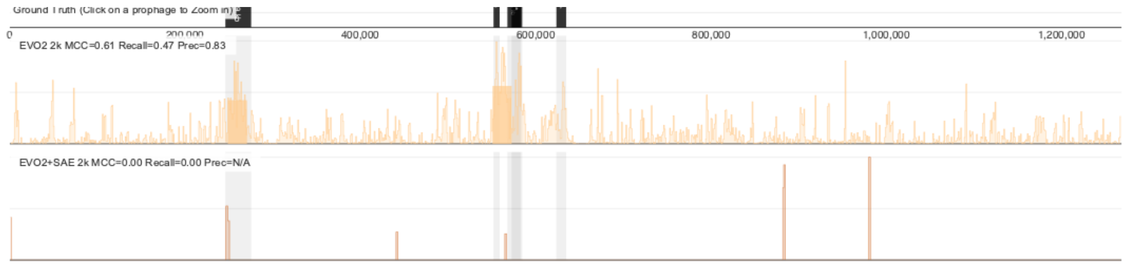


**Figure S18: Genome-wide comparison of EVO2 and EVO2+SAE activations in *Wolbachia* endosymbiont of *Drosophila melanogaster*.** Genome-wide tracks comparing raw EVO2 signal (yellow) and sparse autoencoder activations (orange), with ground truth prophage regions shown in black. EVO2+SAE produces extremely sparse activation across the genome, resulting in no predicted regions (recall = 0.00, precision = N/A), in contrast to EVO2 (recall = 0.47, MCC = 0.61). This example illustrates a case where the SAE feature fails to capture the prophage signal.
